# A molecular plugin rescues GroEL/ES substrates from pre-folding oxidation

**DOI:** 10.1101/2022.05.03.490446

**Authors:** Emile Dupuy, Sander E. Van der Verren, Jiusheng Lin, Mark A. Wilson, Alix Dachsbeck, Felipe Viela, Emmanuelle Latour, Alexandra Gennaris, Didier Vertommen, Yves F. Dufrêne, Bogdan I. Iorga, Camille V. Goemans, Han Remaut, Jean-François Collet

**Author notes:** Address for correspondence: Jean-Francois Collet, Camille Goemans, Han Remaut. These authors contributed equally (Emile Dupuy, Sander E. Van der Verren).

## Abstract

Hsp60 chaperonins and their Hsp10 cofactors assist protein folding in all living cells, constituting the paradigmatic example of molecular chaperones. Despite extensive investigations of their structure and mechanism, crucial questions regarding how these chaperonins promote folding remain unsolved. Here, we report that the bacterial Hsp60 chaperonin GroEL forms a stable, functionally relevant complex with the chaperedoxin CnoX, a protein combining a chaperone and a redox function. Binding of GroES (Hsp10) to GroEL induces CnoX release. Cryo-electron microscopy provided crucial structural information on the GroEL-CnoX complex, showing that CnoX binds GroEL outside the substrate-binding site via a highly conserved C-terminal α-helix. Furthermore, the identification of complexes in which CnoX, bound to GroEL, forms mixed-disulfides with GroEL substrates indicates that CnoX likely functions as a redox quality-control plugin for GroEL. Proteins sharing structural features with CnoX exist in eukaryotes, which suggests that Hsp60 molecular plugins have been conserved through evolution.

## INTRODUCTION

Following synthesis as linear amino acid chains, proteins need to fold to unique three-dimensional (3D) structures to become functional. Seminal work from Anfinsen demonstrated that the information required for a polypeptide to reach its native conformation is contained in its primary sequence (Anfinsen, 1973). For most small proteins, folding to the native state is a spontaneous process that takes less than a few milliseconds (Jahn and Radford, 2005). For larger proteins with multiple domains, however, the path to the native conformation is more tortuous and potentially hazardous. For these proteins, stable intermediates can form, slowing the folding process and potentially leading to aggregation and/or degradation (Ellis, 2001). To deal with this problem, living cells express a network of chaperones that help complex proteins to fold efficiently (Hartl et al., 2011).

The Hsp60 chaperonins are a unique class of chaperones that are essential in all domains of life and prevent unproductive interactions within and between polypeptides using adenosine triphosphate (ATP)-regulated cycles (Hayer-Hartl *et al*., 2016; Horwich and Fenton, 2020). Chaperonins stand out in the proteostasis network as they form a complex tetradecameric structure encompassing a large cylindrical cage consisting of two seven-membered rings stacked back-to-back (**Figure S1A**) (Hendrix, 1979; Hohn et al., 1979). Each Hsp60 subunit consists of an ATP-binding equatorial domain, an intermediate domain, and an apical substrate-binding domain (**Figure S1A**) (Braig et al., 1994). Hsp60 cooperates with Hsp10 (Chandrasekhar et al., 1986), which forms a heptameric dome-like structure (**Figure S1A**) (Hunt et al., 1996). In the presence of nucleotides, Hsp10 associates with the apical domain of Hsp60, binding as a lid covering the ends of the ring and forming a folding chamber (Xu et al., 1997) referred to as the “Anfinsen cage”. Binding of Hsp10 to a substrate-loaded Hsp60 results in displacement of the substrate into the chamber, where it can fold protected from outside interactions (Clare et al., 2012).

The mechanism by which chaperonins assist substrate proteins to navigate the folding landscape to their native state is relatively well understood. Although this is particularly true for *Escherichia coli* GroEL and GroES, its Hsp10 cofactor, several crucial questions remain unsolved. For instance, whether the GroEL-GroES nanomachine actively promotes folding or serves only as a passive folding cage remains controversial (Hayer-Hartl *et al*., 2016). It also remains unknown why some polypeptides are highly dependent on GroEL-GroES for folding whereas homologous proteins with a similar structure fold independently of the chaperonin (Hayer-Hartl *et al*., 2016); thus, further investigation is required to elucidate the sorting signals that recruit substrate proteins to the Hsp60 folding cage. Excitingly, recent results have indicated that the integration of GroEL-GroES in the cellular proteostasis network also needs further exploration. Indeed, whereas GroEL-GroES was thought to largely function in isolation, the identification of CnoX as the first chaperone capable of transferring its substrates to GroEL-GroES for active refolding (Goemans et al., 2018a; Goemans *et al*., 2018b) suggests that functional links between GroEL-GroES and accessory folding factors remain to be discovered. The extreme complexity of the GroEL-GroES molecular machine, its essential role in cell survival, as well as redundancy in the bacterial proteostasis system have slowed progress in the field, highlighting the need for new investigation approaches and experimental strategies.

Here, we sought to explore the details of the newly reported CnoX-GroEL functional relationship (Goemans *et al*., 2018a; Goemans *et al*., 2018b), with the aim of revealing unsuspected features of the GroEL-GroES system. CnoX consists of a redox-active N-terminal thioredoxin domain and a C-terminal tetratricopeptide (TPR) domain (**Figure S1B)** (Lin and Wilson, 2011), a fold often involved in protein–protein interactions. CnoX is a “chaperedoxin,” meaning that it combines a redox-protective function, by which it prevents irreversible oxidation of its substrates, and a holdase chaperone activity, by which it maintains its substrates in a folding-competent state before transferring them to GroEL-GroES for refolding (Goemans *et al*., 2018b). We reasoned that finding the molecular attributes that uniquely allow CnoX to work in concert with GroEL-GroES should lead to new insights into the properties of the GroEL-GroES system.

## RESULTS

### CnoX and GroEL form a stable complex

To start our investigation, we pulled-down CnoX from *E. coli* cellular extracts using specific a-CnoX antibodies. We found that CnoX co-eluted with only one partner (**Figure 1A**), a ∼60-kDa protein identified as GroEL by mass spectrometry (MS), confirming previous results suggesting a direct interaction between the two proteins (Lin and Wilson, 2011). In this experiment, we expressed both CnoX and GroEL from their native locus in cells grown under normal conditions. Exposing the cells to heat shock (42°C) did not lead to an increase in the amount of GroEL that co-eluted with CnoX (**Figure S1C**). We then examined whether the CnoX-GroEL interaction could be reconstituted *in vitro* using purified proteins. *E. coli* CnoX and GroEL were independently overexpressed and purified to near homogeneity (**Figure S1D**). We mixed GroEL and CnoX in a 1:1 molar ratio and found that they co-eluted from both a streptavidin affinity column (**Figure 1B**; a Strep-tag was fused to the N-terminus of CnoX) and a size-exclusion chromatography column (**Figure 1C**). The latter showed the co-eluting GroEL-CnoX complex in an approximately 14:1 molar ratio compared with the 1:1 input ratio. Notably, we also observed that CnoX formed a complex with a GroEL mutant (GroEL_R452A/E461A/S463A/V464A_) known to form a single heptameric ring (**Figure S1E)** (Weissman et al., 1995). Finally, we determined the affinity between the two proteins using fluorescence spectroscopy and fluorescence anisotropy and found that fluorescein-labeled CnoX (FM-CnoX) binds GroEL with a dissociation constant (K_d_) of 310±10 nM (**Figures 1D** and **S1F)**. Using atomic force microscopy (AFM), we measured a specific binding force of 175±75 pN between the two proteins (**Figures S1G** and **S1H**). Thus, we conclude that CnoX physically interacts with GroEL and that the two proteins form a stable complex both *in vitro* and *in vivo*.

**Figure 1.**
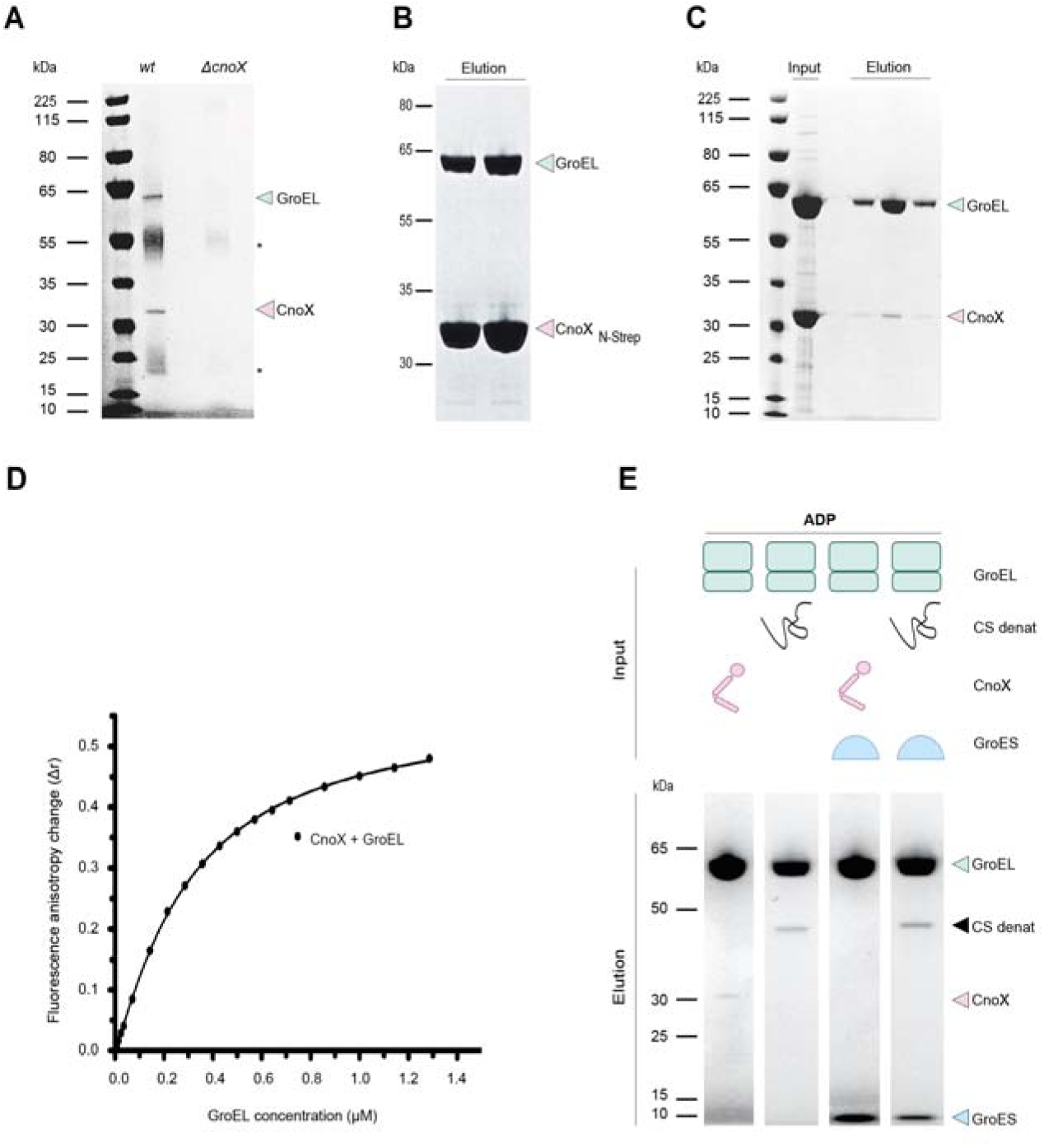
CnoX interacts stably with GroEL. **(A)** GroEL co-elutes with CnoX when CnoX is pulled-down from wild-type cell extracts using α-CnoX antibodies. Both proteins are absent when the experiment is repeated with extracts prepared from the Δ*cnoX* mutant. The image of sodium dodecyl sulphate–polyacrylamide gel electrophoresis (SDS-PAGE), stained with Coomassie blue, is representative of >3 replicates. * indicates the light and heavy chains of the antibodies. **(B)** Purified CnoX_N-Strep_ and GroEL form a complex that can be isolated using streptavidin affinity purification. Two fractions are shown. **(C)** Purified CnoX and GroEL form a complex that can be isolated using size-exclusion chromatography. **(D)** Formation of a complex between FM-CnoX and GroEL can be monitored using fluorescence anisotropy. The non-cooperative model gives an adequate fit to these data, with a K_d_ of 310 nM±10 nM. **(E)** CnoX and unfolded CS co-elute with GroEL from a gel filtration column. Addition of GroES triggers the release of CnoX from GroEL, while CS remains bound to GroEL. Size-exclusion chromatography was performed in the presence of ADP (50 μM), and fractions were analyzed by SDS-PAGE. The results are representative of >3 experiments.

### GroES binding triggers the release of CnoX from GroEL

We next aimed to unravel the interrelationship among CnoX, GroEL, and GroES. GroES reversibly binds GroEL in the presence of nucleotides (Hayer-Hartl *et al*., 2016). The addition of adenosine diphosphate (ADP), which triggers conformational changes in GroEL and primes the ring for GroES binding, had no impact on the GroEL-CnoX complex (purified proteins were mixed in a 14:1 molar ratio) (**Figure 1E**), although the affinity of CnoX for GroEL decreased slightly (K_d_ of ∼350 nM) (**Figure S2A**). Strikingly, however, the subsequent addition of GroES (14[GroEL]:14[GroES]:1[CnoX] molar ratio) triggered the release of CnoX from GroEL (**Figure 1E**), thus indicating a direct or allosteric competition between CnoX and GroES for GroEL binding. We obtained similar results with a non-hydrolysable ATP analogue (**Figure S2B**). Next, titration of a complex between GroEL and FM-CnoX with increasing amounts of GroES resulted in a dose-dependent loss of FM-CnoX, confirming that GroES dissociates CnoX from GroEL (**Figure S2C**). Using a single-site competitive binding model, we calculated a fitted inhibitory constant (K_i_) of 47 nM. Altogether, these results clearly distinguish CnoX from typical GroEL substrates. Indeed, GroEL does not release substrate proteins such as unfolded citrate synthase (CS) upon GroES addition (**Figure 1E**); rather, these proteins become encapsulated inside the GroEL-GroES folding chamber for refolding (Hayer-Hartl *et al*., 2016; Horwich and Fenton, 2020). In the same line, we found that the presence of CnoX does not prevent GroEL from recruiting unfolded CS (**Figure S2D**). Thus, CnoX does not restrict access to the substrate-binding site of GroEL.

### The C-terminal α-helix of CnoX binds GroEL near the site of substrate entry into the cage

Intrigued by these results, we sought to obtain structural information on the CnoX-GroEL interaction using cryoEM. We reconstituted the CnoX-GroEL complex by mixing purified GroEL and CnoX_N-Strep_ (10:1 molar ratio) in the absence of nucleotides. The complex was then affinity-purified (**Figure S3A**) and imaged for single-particle cryoEM analysis (**Figure S3B, S3C** and **Table S1**). Analysis of the two-dimensional (2D) class averages showed the two rings of GroEL stacked back-to-back and revealed the presence of a protruding density on top of the two GroEL rings (**Figures 2A, 2B** and **S3D**). A c7-symmetrical 3D reconstruction resulted in a 3.4-Å electron potential map (**Figure S3E**) showing a density on the GroEL apical domain corresponding to at least five α-helices and allowing an unambiguous rigid body docking with the TPR domain of CnoX (**Figures 2C, 2D, S3F** and **S3G**). The absence of a clearly resolved thioredoxin domain in the CnoX-GroEL complex is consistent with the prior observation of extensive mobility of this domain in the X-ray crystal structure of CnoX alone (Lin and Wilson, 2011). This finding suggests that the thioredoxin domain is highly dynamic, which may be relevant for our proposed model (see below).

**Figure 2.**
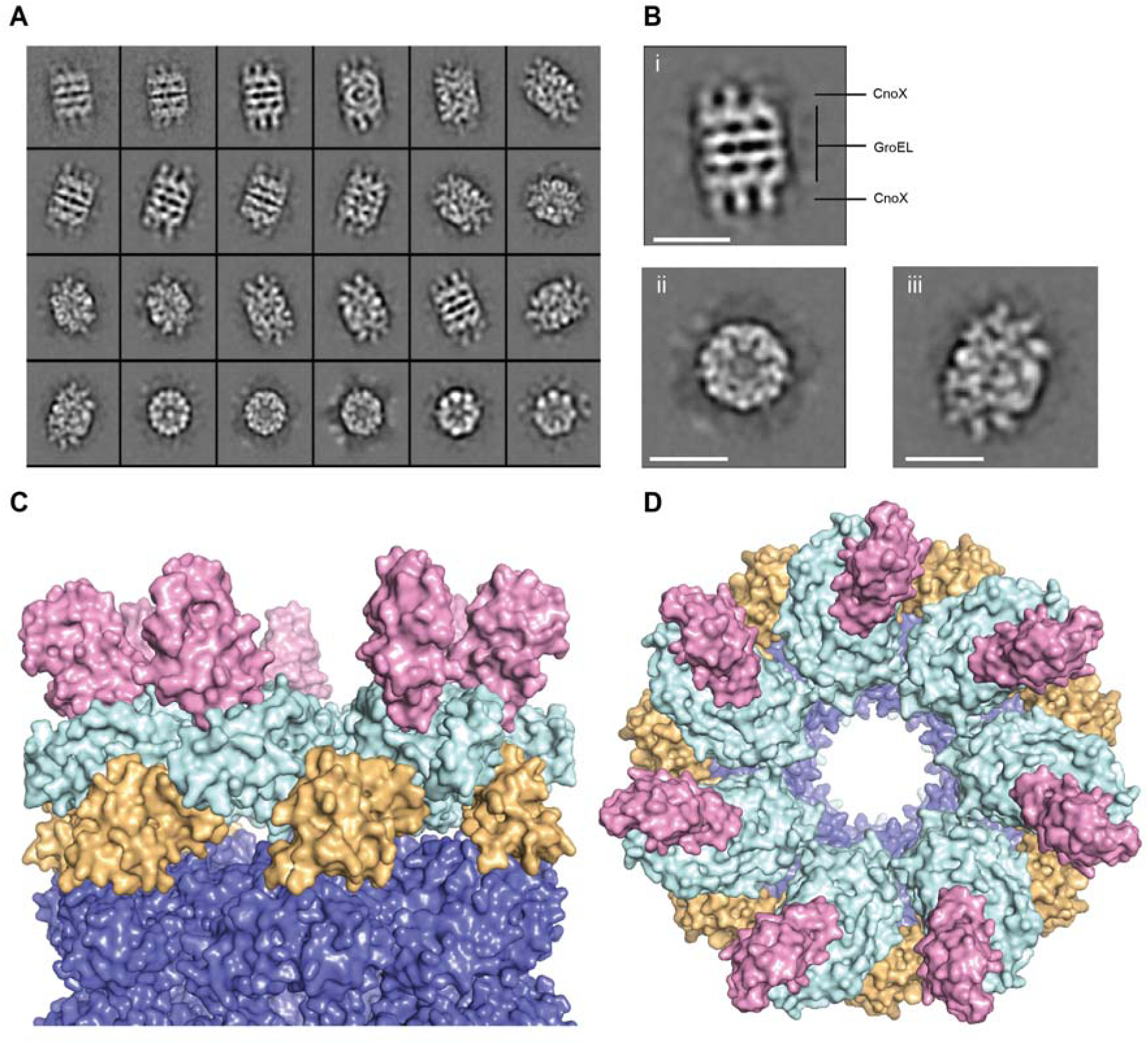
CryoEM shows that the TPR domain of CnoX binds GroEL. **(A-B)** CryoEM 2D class averages of the GroEL-CnoX complex reconstituted *in vitro* at a 10:1 molar ratio (scale bar: 100 Å). **(C-D)** Side and top view of the structure of the GroEL-CnoX complex shown as a solvent-accessible surface. The equatorial, intermediate, and apical domains of GroEL are shown in slate, orange, and light cyan, respectively, and CnoX is shown in pink.

Although the N-terminal thioredoxin domain of CnoX is not visible, the structure provides crucial molecular details regarding the CnoX-GroEL interaction. First, the structure reveals that CnoX binds GroEL via its C-terminal α-helix (**Figure 3A)**; accordingly, a CnoX mutant lacking the last 10 C-terminal residues (CnoX_ΔCter_) is unable to bind GroEL, both *in vivo* (**Figure 3D**) and *in vitro* (**Figure S4A)**. Furthermore, the addition of a His-tag to the C-terminus of CnoX (CnoX_C-His_) prevented CnoX binding to GroEL (**Figures 3D** and **S4A**). Thus, the C-terminal helix of the TPR domain of CnoX functions as a specific GroEL affinity tag that is required for GroEL binding. Interestingly, while the sequence of the TPR domain is diverse among species, the last C-terminal helix is highly conserved (**Figure S4B**) and is structurally and electrostatically distinct from the remainder of the TPR domain (Lin and Wilson, 2011), suggesting that the ability to bind GroEL is widespread and central to CnoX activity. The structure also reveals where CnoX binds to GroEL; the interaction zone, which has a buried surface area of 472 Å^2^ (−4.6 kcal/mol; PDBePISA (Brinker et al.)) and encompasses residues D224, K286–M307, K311, D316, R345, and Q348 (**Figures 3B** and **3C**), corresponds to a shallow surface cleft formed by helices J and K in the apical domain of GroEL. This region does not overlap with the substrate-binding site of GroEL in helices H and I (Hayer-Hartl *et al*., 2016; Horwich and Fenton, 2020), as also corroborated by the above results (**Figure S2D**). At least five potential H-bond or electrostatic interactions stabilize the contacts between CnoX and GroEL (R255–E304, R277–G298, R277–T299, Y284–E304, and Y284–R345, listed as CnoX–GroEL), as well as a hydrophobic interaction by CnoX residues L279, Y280, and L283 and GroEL residues V300, I305, and M307 (**Figures 3B** and **3C**). Accordingly, introducing a set of mutations in the interaction interface disrupted the GroEL-CnoX interaction (**Figure 3E)**. GroEL is a highly dynamic protein that undergoes substantial conformational rearrangements depending on the binding of a nucleotide, position in the folding pathway, or binding of GroES (Clare *et al*., 2012). Comparison of our structure with the different conformational states of GroEL shows that the rings of GroEL are in a conformation corresponding to that of the nucleotide-free protein (**Figure S5**), as expected. Our findings also indicate that the CnoX-binding paratope remains fully accessible in all conformations, except when GroES is bound (**Figure S5**). The persistence of the CnoX-binding site in various conformations of GroEL is consistent with the ability of CnoX to bind to GroEL irrespective of the presence of a nucleotide (**Figures 1B, 1C, 1E** and **S2B**). Available structures also show a large conformational rotation of the GroEL apical domain in the GroEL-GroES complex. Although the GroES-binding site does not directly overlap with that of CnoX, the conformation of the apical domain results in a steric occlusion of the CnoX-binding paratope (**Figure S5)**, providing a molecular explanation to our finding that GroES docking onto GroEL is incompatible with CnoX binding (**Figure 1E** and **S2B**).

**Figure 3.**
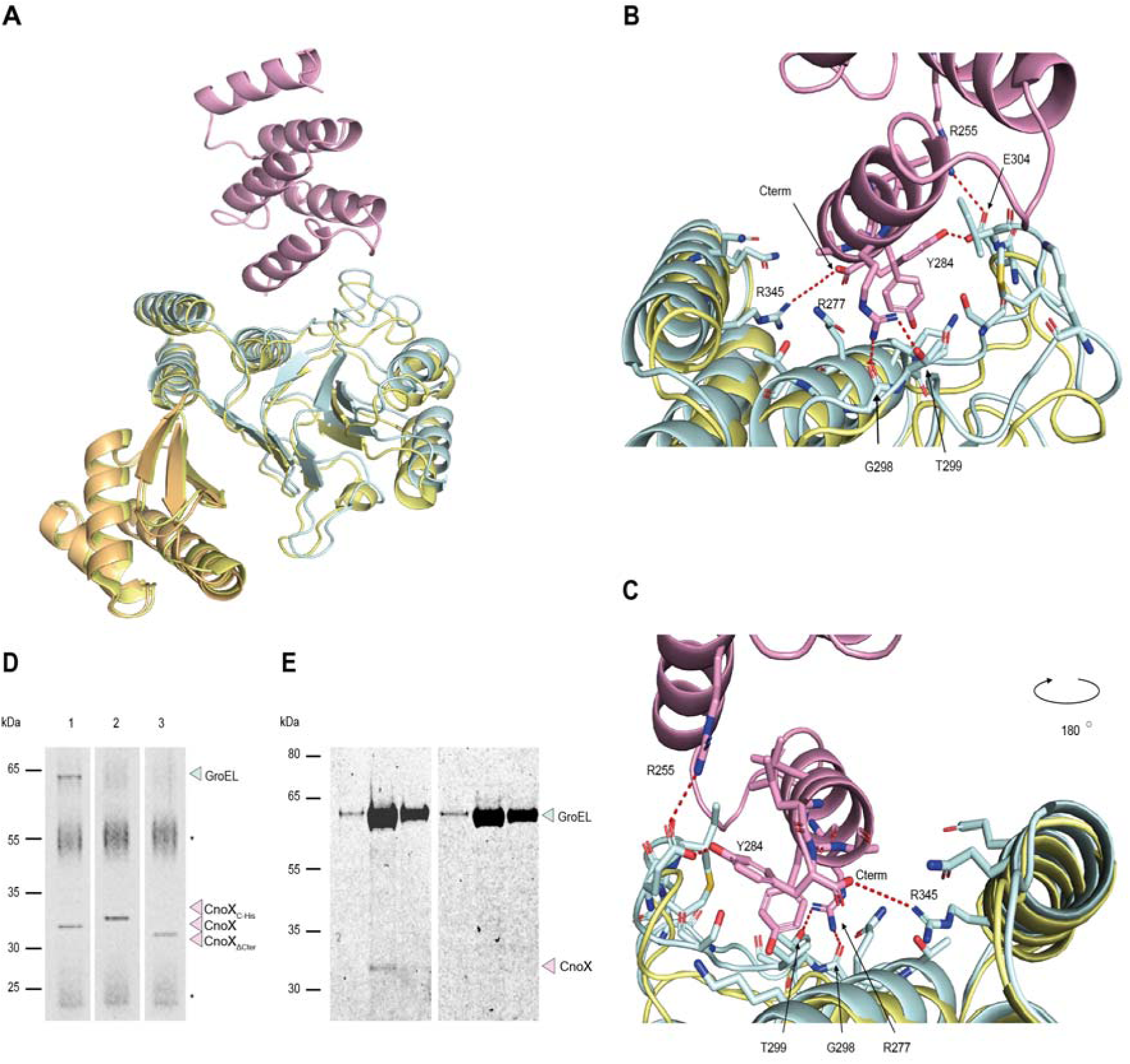
The C-terminal α-helix of CnoX binds a shallow cleft in the apical domain of GroEL. **(A)** Ribbon representation of a single GroEL-CnoX protomer. CnoX binds GroEL via its C-terminal α-helix. The intermediate and apical domains of GroEL are shown in orange and light cyan, respectively. CnoX is shown in pink. For comparison, the GroEL-CnoX structure is shown superimposed on the structure of T state GroEL (yellow; PDB: 1grl). **(B-C)** Close-up views of the GroEL-CnoX binding interface. CnoX binds GroEL through the following H-bond and electrostatic interactions (CnoX–GroEL): R255–E304, R277–G298, R277–T299, Y284–E304, and Y284 C-term–R345. For comparison, the GroEL-CnoX structure is shown superimposed on the structure of T state GroEL (yellow; PDB: 1grl). **(D)** GroEL co-elutes with CnoX (lane 1) but not with CnoX_C-His_ (lane 2) or CnoX_ΔC-ter_ (lane 3) when CnoX is pulled-down from cell extracts using α-CnoX antibodies. In these experiments, CnoX, CnoX_ΔC-ter_, and CnoX_C-His_ were expressed in Δ*cnoX* cells. The SDS-PAGE gel, stained with Coomassie blue, is representative of >3 replicates. * indicates the light and heavy chains of the antibodies. **(E)** GroEL^§^, a GroEL variant with mutations in the CnoX-binding site (G298A/T299L/V300K/E304L/I305K/M307K/R345L), does not elute together with CnoX from a size-exclusion chromatography column (right), in contrast to wild-type GroEL (left). Three consecutive elution fractions are shown for each chromatography.

### CnoX forms mixed-disulfides with obligate GroEL substrates when bound to GroEL

We next aimed to gain insight into the physiological relevance of the CnoX-GroEL complex *in vivo*. GroEL-GroES substrates often need minutes to fold after leaving the ribosome (Ewalt et al., 1997), which raises a question regarding how their amino acids are protected from oxidative damage before reaching their native state. This question is particularly relevant for cysteine residues, which are highly sensitive to oxidation by the molecular oxidants that are present in cells even in the absence of stress (Ezraty et al., 2017; Imlay, 2008). Indeed, the thiol side chain of a cysteine is readily oxidized to a sulfenic acid (−SOH), an unstable derivative that can react with another cysteine in the vicinity to form a disulfide or that can be irreversibly oxidized to sulfinic and sulfonic acids. Similar to Anfinsen’s experiments showing that noncanonical disulfide pairing thwarts *in vitro* protein folding, one can expect the GroEL chaperonin to require its substrates’ cysteines to be reduced for proper folding. CnoX stands out in the proteostasis network in that it combines a chaperone and a redox-protective function (Goemans *et al*., 2018b); therefore, CnoX may bind GroEL to function as a redox rescue mechanism for slow-folding GroEL-GroES substrates.

By performing additional pull-down experiments, we obtained a crucial result shedding light onto the function of CnoX. When GroEL is pulled-down from cellular extracts, it co-elutes with CnoX, as expected. Intriguingly, we found that high-molecular-weight complexes involving CnoX are also pulled-down (**Figure 4A**). When a reducing agent was added, these complexes disappeared, indicating that they correspond to mixed disulfides comprising CnoX and unknown proteins. Accordingly, we did not detect high-molecular-weight complexes when the experiment was repeated with a CnoX mutant lacking the two cysteine residues (CnoX_no_cys_; **Figure 4A**). We identified the proteins involved in the mixed disulfides using MS (**Table S2**); excitingly, we found that these proteins include several obligate GroEL substrates (**Figure 4B** and **Table S2**). Thus, we conclude that CnoX forms mixed disulfides with obligate GroEL substrates when bound to GroEL in the cell.

**Figure 4.**
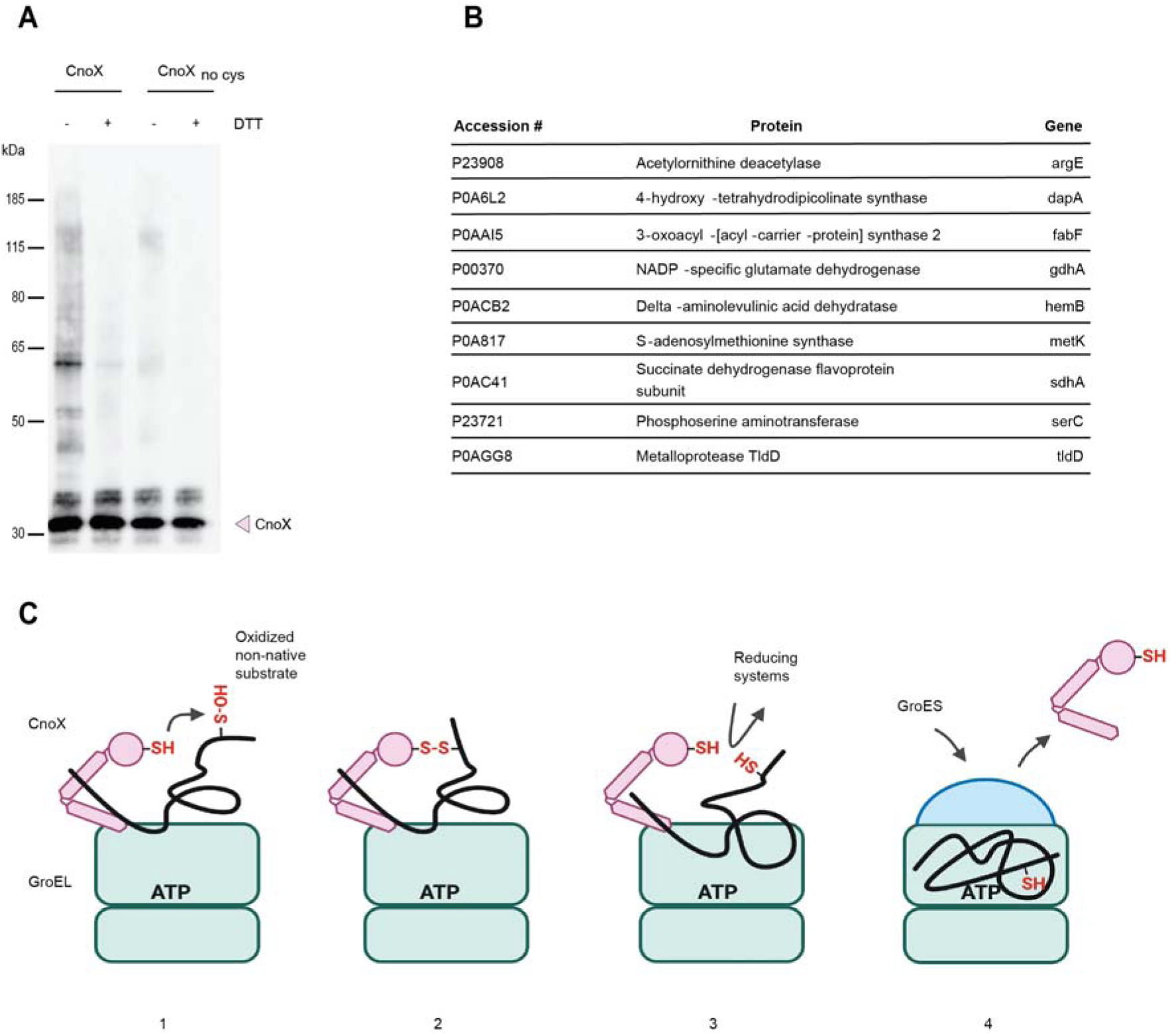
CnoX functions as a molecular plugin to rescue GroEL substrates from oxidative damage. **(A)** CnoX co-elutes with GroEL when the chaperonin is pulled-down from wild-type cell extracts using specific antibodies. High-molecular-weight complexes corresponding to dithiothreitol (DTT)-sensitive mixed disulfides are detected by α-CnoX antibodies. These complexes are not detected when the experiment is repeated using extracts from cells expressing a CnoX mutant lacking the two cysteine residues, CnoX_no_cys_. **(B)** Obligate GroEL substrates trapped in mixed-disulfide complexes with CnoX and pulled-down using α-GroEL antibodies were identified using liquid chromatography with tandem MS (LC-MS/MS). **(C)** Model: 1. CnoX forms a stable complex with GroEL via its C-terminal α-helix in a nucleotide-independent manner. Positioned on the apical domain of GroEL, CnoX interacts with incoming substrates for GroEL, acting as a redox quality-control plugin 2. If the substrate that reaches GroEL for folding presents oxidized cysteine residues (to a sulfenic acid or in a disulfide bond), CnoX reacts with the substrate via the cysteines of its thioredoxin domain, and a mixed disulfide is formed. 3. Cytoplasmic reducing pathways then reduce the mixed disulfide, releasing the substrate in a reduced, folding-competent state. 4. GroES binding then triggers CnoX release from GroEL and encapsulation of the substrate within the folding cage for folding.

### CnoX functions as a molecular plugin providing redox quality-control for GroEL substrates

Altogether, our results suggest the following model (**Figure 4C**). Regardless of stress, CnoX binds GroEL via its highly conserved C-terminal α-helix in a nucleotide-independent manner. The CnoX-binding interface on GroEL does not overlap with the substrate-binding site. If the substrate that reaches GroEL for folding presents oxidized cysteine residues (to a sulfenic acid or in a disulfide bond), CnoX reacts with the substrate via the cysteines of its thioredoxin domain, resulting in the formation of a mixed disulfide. Cytoplasmic reducing pathways then reduce the mixed disulfide, releasing the substrate in a reduced, folding-competent state. The binding of GroES to GroEL induces conformational changes in the chaperonin and occludes the CnoX-binding site, triggering CnoX release from GroEL and encapsulation of the substrate within the folding cage for folding. Thus, we propose that CnoX functions as a molecular plugin that provides redox quality-control for GroEL substrates. Our model is compatible with both the binding of CnoX to unfolded oxidized client proteins in solution followed by delivery to the GroEL chaperonin and the surveillance performed by CnoX to identify erroneously oxidized client proteins that may become stuck at the substrate entrance to the Anfinsen cage of GroEL.

## DISCUSSION

Investigations of Hsp60 chaperonins started in the 1970s (Horwich and Fenton, 2020), when researchers described mutations that blocked phage head assembly in *groE* and discovered the tetradecameric structure of GroEL, the archetypical member of the Hsp60 family, using electron microscopy (EM). Since then, a large body of studies has examined the mechanistic and structural properties of Hsp60 proteins and their Hsp10 co-chaperones, not only in bacteria but also in chloroplasts and mitochondria (Horwich and Fenton, 2020). This impressive amount of work has rendered chaperonins a textbook example of folding systems. In the current study, the identification of CnoX as a quality-control protein that physically interacts with GroEL-GroES for optimal folding further widens this field of investigation by uncovering a novel, unsuspected feature of Hsp60s. Additional questions remain unsolved and will be the subject of future research. For instance, the biologically active stoichiometry of the CnoX-GroEL complex warrants careful investigation, as well as the specific role of the cytoplasmic reducing pathways in the reduction and release of mixed disulfides. Future work must also establish the location of the N-terminal thioredoxin domain when CnoX is bound to GroEL. Our results show that CnoX forms mixed disulfides with GroEL substrates while being bound to GroEL, but future research will elucidate whether CnoX also functions as a tugboat to locate endangered GroEL substrates in the cytoplasm and escort them to the chaperonin. Finally, it will be important to determine whether similar proteins with a redox quality-control function exist in other organisms, including eukaryotes. The facts that *E. coli* CnoX stably interacts with human mitochondrial Hsp60 (**Figure S6A**) and that proteins sharing structural features with CnoX exist in eukaryotes (**Figure S6B, S6C** and **S6D**) support this idea. Along the same line, it is tempting to speculate that living cells could also contain Hsp60 molecular “plugins” with specific, redox-independent functions yet to be discovered.

## SUPPLEMENTAL INFORMATION

Supplemental information (Methods, Figures S1 to S6, Tables S1 to S5) can be found online at …

## ACKNOWLEDGEMENTS

We thank Asma Boujtat and Gaetan Herinckx for technical help. We are indebted to Dr. Michael Deghelt, Dr. Seung-Hyun Cho, and Dr. Pauline Leverrier for helpful suggestions and discussions and for providing comments on the manuscript. M.A.W. is supported by National Institutes of Health grant R01GM139978. We thank Dr. Tommi White and Dr. Javier Seravalli for assistance with the GroEL-CnoX quantitative interaction studies and Dr. Aron Fenton for discussions about cooperative protein binding. We thank staff at VIB-VUB facility for Bio Electron Cryogenic Microscopy (BECM) for assistance in data collection. This work was funded by the Fonds de la Recherche Scientifique (FNRS) grant agreements WELBIO-CR-2015A-03 and WELBIO-CR-2019C-03, the EOS Excellence in Research Program of the FWO and FRS-FNRS (G0G0818N), the Fédération Wallonie-Bruxelles (ARC 17/22-087), the GENCI-IDRIS (2021-A0100711524), the Flanders Research Foundation Hercules grant (G0H5916N), the Flanders Research Foundation PhD fellowship programme, and the Flanders Institute for Biotechnology – VIB.

## DATA AVAILABILITY

Coordinates and the electron potential maps for the GroEL:CnoX cryoEM structure have been deposited in the PDB and EMDB under accession codes 7YWY and EMD-14352, respectively. All other data generated or analyzed during this study are included in this published article and its supplementary information file.

## AUTHOR CONTRIBUTIONS

Writing: JFC, SVdV, ED, HR, and CVG. Conceptualization: CVG, ED, HR, and JFC. Investigation, strain construction, construct cloning: ED, CVG, AD, AG, SVdV, JL, MAW, EL, YFD, and FV. Interactive molecular-dynamics flexible fitting of the cryoEM model: BII. Mass spectrometry: DV. Data analysis and interpretation: ED, SVdV, CVG, HR, JL, MAW, YFD, FV and JFC. All authors discussed the results and commented on the manuscript.

