## Supplementary Information for "A molecular plugin rescues GroEL/ES substrates from pre-folding oxidation"

**Methods**

**Bacterial strains and plasmids**

The bacterial strains used in this study are listed in **Table S3**. We prepared the *cnoX* deletion mutant by transferring the corresponding allele (*ybbN*::kan^R^) from the Keio collection (Baba et al., 2006) into MG1655 by P1 phage transduction, with verification by polymerase chain reaction (PCR). To excise the kanamycin-resistance cassette, we used pCP20 (Cherepanov and Wackernagel, 1995). Unless otherwise indicated, cells grew in Lysogeny broth (LB) at 37°C, and when necessary, we supplemented the growth media with ampicillin (100–200 µg/mL), chloramphenicol (100 µg/mL), or kanamycin (50 µg/mL).

**Table S4** shows the plasmids used in this study. The primers used for their construction are shown in **Table S5**. To obtain high expression levels of wild-type CnoX, CnoX_C-His_, GroEL, and GroES, we cloned the corresponding genes into the high copy vector pET22b(+) (Novagen). DNA encoding CnoX_N-Strep_ was cloned into the medium copy vector pACYCDuet-1. To prepare an expression vector for *cnoX* without the last 10 C-terminal residues (CnoX_∆Cter_), we amplified the corresponding sequence from *E. coli* genomic DNA and ligated the sequence into pET22b(+). The CnoX and GroEL substitution variants used in this study were generated by site-directed mutagenesis, except for GroEL_G298A/T299L/V300K/E304L/I305K/M307K/R345L_, which was generated using Gibson assembly. We generated the CnoX_N-link/C38A/C63A_ variant using site-directed mutagenesis by inserting the coding sequences of Met, Ala, Cys, Ala, and Gly residues at the N-terminal extremity. To prepare an expression vector for human mitochondrial Hsp60, we amplified the corresponding sequence from the MAC_C_CH60 vector (Addgene) without the mitochondrial targeting sequence (MTS), which was replaced by the coding sequences of Met, Gly, and Ser residues, and cloned the sequence into pET22b(+) using Gibson assembly.

**Expression and purification of CnoX, CnoX_∆Cter_, CnoX_N-Strep_, CnoX_C-His_, and CnoX_N-link/C38A/C63A_**

We utilized *E. coli* BL21 (DE3) carrying pET22b_cnoX, pET22b_cnoX_∆Cter_, or pET22b_cnoX_N-link/C38A/C63A_ to overexpress wild-type CnoX, CnoX_∆cter_, and CnoX_N-link/C38A/C63A_, respectively. Cells grew at 37°C until an optical density at 600 nm (OD_600_) of 0.5 was reached, and then, we added 1 mM isopropyl β-D-1-thiogalactopyranoside (IPTG) for 3 h to induce protein expression. Cells were pelleted, resuspended in 50 mM Tris-HCl pH 8, and disrupted with a French press. After centrifugation for 30 min at 40,000*g* at 4°C, the supernatant was filtered through 0.45-μm filters and loaded onto a 5-mL Q-Sepharose HP column (GE Healthcare). We eluted the proteins with a 0%–50% gradient of 1 M NaCl in 50 mM Tris-HCl pH 8. Proteins underwent a second step of purification via a HiLoad 16/60 Superdex 75 gel filtration system (GE Healthcare) and were eluted in 10 mM hydroxyethyl piperazineethanesulfonic (HEPES)-KOH pH 8, 100 mM NaCl.

We employed *E. coli* BL21 (DE3) carrying pET22b-cnoX_C-His_ to overexpress CnoX fused to a C-terminal His-tag (CnoX_C-His_). Cells grew at 37°C until an OD_600_ of 0.5 was reached, and we added 1 mM IPTG for 3 h to induce protein expression. Cells were pelleted, resuspended in buffer A (NaPi 50 mM pH 8, 300 mM NaCl), and lysed with a French press. After centrifugation for 30 min at 40,000*g* at 4°C, the supernatant was filtered through 0.45-μm filters and loaded on Ni-nitriloacetic acid (NTA) agarose beads (5 mL; IBA Lifesciences) previously equilibrated with buffer A. After washing the resin with buffer A supplemented with 20 mM imidazole, we eluted the proteins with buffer A supplemented with 300 mM imidazole. As a final purification step, we performed size-exclusion chromatography using a HiLoad 16/60 Superdex 75 column (GE Healthcare) with 10 mM HEPES pH 7, 100 mM NaCl.

*E. coli* BL21 (DE3) carrying pACYCDuet-1-cnoX_N-Strep_ was used to express CnoX fused to an N-terminal Strep-tag (CnoX_N-Strep_). Cells grew at 37°C until an OD_600_ of 0.5 was reached, and we added 1 mM IPTG for 3 h to induce protein expression. Cells were pelleted, resuspended in buffer B (50 mM Tris-HCl pH 8, 150 mM NaCl), and lysed with a French press. After centrifugation for 30 min at 40,000*g* at 4°C, the supernatant was filtered through 0.45-μm filters and loaded onto a 5-mL Strep-Tactin column (IBA Lifesciences) previously equilibrated in buffer B. After a washing step with buffer B, we performed elution with buffer B supplemented with 2.5 mM D-desthiobiotin. We then purified the sample using size-exclusion chromatography on a HiLoad 10/300 Superdex 200 column (GE Healthcare) using buffer B supplemented with 1 mM ethylenediaminetetraacetic acid (EDTA).

**Expression and purification of GroEL, GroEL variants, and GroES**

We used *E. coli* BL21 (DE3) cells carrying pET22b-groL to overexpress GroEL. Cells grew at 37°C until an OD_600_ of 0.5 was reached, and we added 1 mM IPTG for 3 h to induce protein expression. Cells were pelleted and resuspended in buffer C containing 50 mM Tris-HCl pH 7.4, 1 mM EDTA, and 1 mM DTT. After centrifugation for 30 min at 40,000*g* at 4°C, the supernatant was filtered through 0.45-μm filters, loaded onto a 5-mL DEAE-Sepharose HP column (GE Healthcare), and eluted in a gradient (0%–50%) of 1 M NaCl in buffer C. The second purification step required gel filtration using a HiLoad S200 16/600 column (GE Healthcare) and elution buffer containing 10 mM HEPES-KOH pH 8, 100 mM NaCl, and 1 mM DTT. Finally, we loaded the protein onto a 5-mL Q-Sepharose HP column (GE Healthcare) using a gradient (0%–50%) of 1 M NaCl in buffer C. GroEL_D490C_, GroEL_R452A/E461A/S463A/V464A_, and GroEL_G298A/T299L/V300K/E304L/I305K/M307K/R345L_ (GroEL^§^) were purified in a similar manner.

We utilized *E. coli* BL21 (DE3) carrying pET22b-groS to overexpress GroES. Cells were grown at 37°C. When an OD_600_ of 0.5 was reached, we added 1 mM IPTG for 3 h to induce protein expression. Cells were pelleted and resuspended in buffer D (30 mM Tris-HCl pH 7.5, 10 mM NaCl, 1 mM EDTA). After centrifugation for 30 min at 40,000*g* at 4°C, the supernatant was filtered (0.45-μm filters), loaded onto a 5-mL DEAE-Sepharose HP column (GE Healthcare), and eluted in a 0%–50% gradient of 1 M NaCl in buffer D. For the second purification step, we loaded the sample onto a 5-mL Q-Sepharose HP column (GE Healthcare), and proteins were eluted with a 0%–50% gradient of 1 M NaCl in 20 mM imidazole pH 5.8. Finally, we loaded the sample onto a gel filtration column (HiLoad S200 26/60 column [GE Healthcare]) equilibrated in buffer D.

For all proteins, we verified the purity via SDS-PAGE with Coomassie staining and concentrated samples using a Vivaspin Turbo (Sartorius) with a 5-kDa molecular-weight cutoff. We assessed protein concentrations by measuring the absorbance at 280 nm using a Varian Cary 50 UV-Vis spectrophotometer.

**Reconstitution of the GroEL-CnoX_N-Strep_ complex**

To reconstitute GroEL-CnoX_N-Strep_, we mixed GroEL (150 µM) in excess (10:1) or with equimolar amounts of CnoX_N-Strep_ in 500 µL of buffer A. After 15 min at room temperature, the mixture was loaded onto a 1-mL Strep-Tactin column (IBA Lifesciences) previously equilibrated in buffer A. After a washing step with buffer A, we performed elution with buffer A supplemented with 2.5 mM D-desthiobiotin.

***In vivo* pull-down of CnoX**

*E. coli* MG1655 wild-type (wt), *∆cnoX*, and *E. coli* MG1655 cells carrying pET22b-cnoX, pET22b-cnoX_∆Cter_, or pET22b-cnoX_C-His_ grew in LB (25 mL) at 37°C in a shaking incubator until the mid-log phase was reached. The cells were then harvested, resuspended in 1 mL of buffer E (50 mM Tris-HCl pH 8, 150 mM NaCl, 1 mM EDTA), and sonicated. After a 5-min centrifugation at 16,000*g* at 4°C, we added 5 µL of undiluted rabbit anti-CnoX antibody and 50 µL of protein A/G magnetic beads (Pierce) to the supernatant to immunoprecipitate CnoX; then, we incubated the samples for 30 min on a wheel at room temperature. After three washes with 500 µL of buffer E, CnoX was eluted with 20 µL of 100 mM glycine pH 2.5. We neutralized the samples with 2 µL of 1.5 M Tris-HCl pH 8 before SDS-PAGE analysis and Coomassie staining.

***In vitro* pull-down of GroEL**

We mixed GroEL (5 µM) with equimolar amounts of either CnoX, CnoX_∆Cter_, CnoX_C-His_, or CnoX and unfolded CS in 100 µL of buffer E. After 30 min at room temperature, GroEL was immunoprecipitated by adding 5 µL of undiluted rabbit a-GroEL antibody and 50 µL of protein A/G magnetic beads (Pierce); samples were incubated for 30 min on a wheel at room temperature. After three washes with buffer E, GroEL was eluted with 20 µL of 100 mM glycine, pH 2.5. We then neutralized the samples with 2 µL of 1.5 M Tris-HCl pH 8 before SDS-PAGE analysis and Coomassie staining.

***In vitro* pull-down of CnoX**

CnoX, CnoX_∆Cter_, or CnoX_C-His_ (5 µM) was mixed with equimolar amounts of GroEL in 100 µL of buffer E. After 30 min at room temperature, we immunoprecipitated CnoX by adding 5 µL of undiluted rabbit a-CnoX antibody and 50 µL of protein A/G magnetic beads (Pierce); then, we incubated the samples for 30 min on a wheel at room temperature. After three washes with buffer E, CnoX was eluted with 20 µL of 100 mM glycine pH 2.5. We neutralized the samples with 2 µL of 1.5 M Tris-HCl pH 8 before SDS-PAGE analysis and Coomassie staining.

**Size-exclusion chromatography analysis**

To reconstitute GroEL-CnoX, we mixed both proteins in a 1:1 molar ratio (a 14:1 molar ratio was used with GroEL^§^). After 15 min at room temperature, the sample was loaded onto a Superdex S200 10/300 column equilibrated with buffer E. The fractions corresponding to absorbance peaks at 280 nm were collected and analyzed by gradient (4%–12%) SDS-PAGE.

To determine the impact of GroES addition on the GroEL-CnoX complex, we mixed GroEL (45 µM) with CnoX at a molar ratio of 14:1 in 1 mL of buffer F (50 mM Tris-HCl pH 8, 150 mM KCl, 10 mM MgCl_2_, 1 mM DTT, 1 mM EDTA) supplemented with either ADP (1 mM) or AMP-PNP (1 mM). After 15 min at room temperature, GroES was added at a molar ratio of 14:1:14 (GroEL:CnoX:GroES). After 15 min, we loaded the sample onto a Superdex S200 10/300 column equilibrated with buffer A supplemented with ADP (50 µM) or AMP-PNP (50 µM). The column ran at a flow rate of 0.5 mL/min with 1-mL fractions using an AKTA Purifier (Cytiva). Fractions corresponding to absorbance peaks at 280 nm were collected and analyzed by gradient (4%–12%) SDS-PAGE.

To investigate the binding of CS on GroEL, CS (82 µM, Sigma-Aldrich) was unfolded by dilution in 4 M guanidinium chloride and incubated at room temperature for 2 h. To assay binding on GroEL, we mixed GroEL (45 µM) with unfolded CS (CS_denat_) at a molar ratio of 14:1 GroEL:CS_denat_ in 1 mL of buffer F supplemented with 1 mM ADP. After 15 min at room temperature, GroES was added at a molar ratio of 14:1:14 GroEL:CSdenat:GroES; after another 15 min, we loaded the mixture onto a Superdex S200 10/300 column equilibrated with buffer A supplemented with 50 µM ADP. The column ran at a flow rate of 0.5 mL/min using an AKTA Purifier, and 1-mL fractions were collected. The fractions corresponding to absorbance peaks at 280 nm were collected and analyzed by gradient (4%–12%) SDS-PAGE.

**Purification of GroEL, GroES, and CnoX for quantitative binding studies**

CnoX and GroES were expressed and purified as previously described (Lin and Wilson, 2011). Briefly, proteins were purified by Ni^2+^-NTA metal affinity chromatography, had their N-terminal hexahistidine tags removed by thrombin cleavage, and were passed again over Ni^2+^-NTA resin. These proteins were then collected in the flow-through, dialyzed into storage buffer (25 mM HEPES pH 7.5, 100 mM KCl), snap-frozen in 50- to 100-μL aliquots in liquid nitrogen, and stored at -80°C. We verified tag removal by electrospray MS. We induced GroEL expression from pET15b as an untagged protein by removing the sequence between the “CC” of the NcoI site and the “ATG” of the NdeI site using mutagenic PCR primers (see **Table S5**). GroEL lacking any tag or vector-derived amino acids was expressed by 0.5 mM IPTG induction, and cells were harvested by centrifugation as described. After sonication and centrifugation at 12000*g*, clarified lysate was loaded onto a Macro-Prep High Q resin (Bio-Rad) column, and proteins were eluted over a 0.1–1.1 M NaCl gradient in 25 mM HEPES pH 7.5. The fractions ran on SDS-PAGE gels, and those enriched in GroEL were pooled. We precipitated the crude GroEL fraction by gradually adding finely ground (NH_4_)_2_SO_4_ powder to 75% saturation at room temperature and centrifuged the fraction at 20,000*g* for 45 min. The pellet was then dissolved in storage buffer (25 mM HEPES pH 7.5, 100 mM KCl) and dialyzed against storage buffer overnight. GroEL was applied to a Sephacryl S-300 HR gel filtration column and eluted near the void volume due to the large size of the GroEL oligomer. We identified the fractions containing GroEL by SDS-PAGE and then pooled and dialyzed the fractions against 50 mM Tris-HCl pH 7.5, 1 mM EDTA, 30% methanol (Tris-methanol buffer). The pooled GroEL fractions were loaded onto a UNO Q12 anion exchange column (Bio-Rad) and eluted with a gradient of 0–1 M NaCl in Tris-methanol buffer. We identified GroEL-containing fractions by SDS-PAGE, and those lacking any visible contaminants were pooled, dialyzed into storage buffer, concentrated by centrifugation with a 10-kDa molecular-weight cutoff concentrator, snap-frozen in 50- to 100-μL aliquots in liquid nitrogen, and stored at -80°C.

**Fluorescent labeling of CnoX**

We fluorescently labeled CnoX using fluorescein-5-maleimide (FM; Thermo Fisher Scientific) for binding studies. The maleimide moiety of FM is cysteine-reactive and is therefore expected to label the N-terminal thioredoxin-like domain of CnoX, which contains the only two cysteine residues in the protein, Cys38 and Cys63. A 25-fold molar excess of FM to CnoX in 20 mM sodium phosphate buffer pH 7.2, 150 mM NaCl, and 5 mM EDTA was incubated for 4 h at room temperature in the dark. We then dialyzed the sample using 3.5-kDa-cutoff regenerated cellulose tubing against storage buffer overnight at 4°C to remove any unreacted FAM.

We determined the stoichiometry of FM-CnoX labeling via the ratio of absorbance at 495 nm and 280 nm.

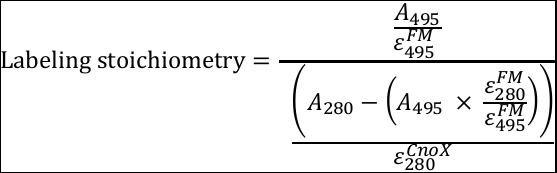

The calculated ε_280_ for CnoX is 23,000 M^-1^ cm^-1^ (ExPASY), the ε_495_ of FAM is 75,000 M^-1^cm^-1^, and the ratio of FM extinction coefficients at 280 and 495 nm ($\frac{\varepsilon_{280}^{FM}}{\varepsilon_{495}^{FM}}$; sometimes called the correction factor) is 0.3. The labeling stoichiometry is 2.2 FAM per CnoX molecule, as expected for the two cysteine residues in the N-terminal thioredoxin domain. FM-CnoX was snap-frozen in liquid nitrogen and stored at -80°C.

**Measurement of the CnoX-GroEL binding affinity**

We performed fluorescence measurements on a Cary Eclipse fluorescence spectrophotometer (Varian) at room temperature (~22°C). We measured fluorescence emission spectra between 500 and 560 nm with an excitation wavelength of 495 nm. The excitation and emission slits were set to 5 nm, and spectra were recorded at a scan rate of 10 nm/s. To measure the binding of GroEL to FM-CnoX, we titrated 119 μM GroEL into 0.5 μM FM-CnoX in a total volume of 0.5 mL storage buffer. Final GroEL concentrations ranged from 0.035 μM to 1 μM. We calculated the normalized fluorescence ratio (D) as:

$$D=\frac{F_{i}}{F_{0}}-1$$

where Fi is the fluorescence emission of FM-CnoX at peak (typically 520 nm) measured after the addition of GroEL and F0 is the fluorescence emission at peak measured for FM-CnoX alone.

To complement the fluorescence emission measurements, we performed a fluorescence anisotropy binding assay using a Jasco J-815 CD spectrophotometer at 20°C. We employed an excitation wavelength of 495 nm and a 540-nm emission filter. Measurements were collected using a cell with a 1-cm path length at a scan rate of 0.833 nm/s from 450 to 550 nm. In a total volume of 2 mL of storage buffer, we titrated 0.5 μM FM-CnoX with 119 μM GroEL to final concentrations ranging from 0.036 μM to 1 μM. The final volume change after GroEL titration was less than 1%. We measured the fluorescence anisotropy for each titration point 2 min after the addition of GroEL. The data were averaged from 3–4 measurements. Binding data were plotted and fitted in Prism (GraphPad) using nonlinear least-squares minimization. We utilized a single-site binding model with positive cooperativity for the CnoX-GroEL interactions and a single-site competition model with a logarithmic competitor concentration for the GroES competition experiment. No linearization of the data was performed.

**Influence of nucleotides on the CnoX-GroEL and GroES interaction**

To examine the effect of ATP or ADP on GroEL-CnoX binding, we repeated the experiment described in the previous section using 0.5 μM FM-CnoX and GroEL in storage buffer with 2 mM ATP (Alfa Aesar) or 2 mM ADP (Sigma) and 10 mM MgCl_2_. For the GroES competition experiment, 0.5 μM FM-CnoX was incubated with 0.5 μM GroEL, with or without 2 mM ATP in storage buffer and then titrated with GroES to final concentrations ranging from 0.035 μM to 2 μM. We measured the fluorescence emission at peak 2 min after addition of the titrated protein at room temperature. In all titrations, the final volume change was less than 1%; hence, no dilution correction was performed. Data were averaged from 3–4 measurements.

***In vivo* trapping of GroEL-CnoX-substrate tripartite complexes**

To identify proteins trapped in a mixed disulfide with CnoX when CnoX is bound to GroEL, we immunoprecipitated GroEL from extracts prepared from *E. coli* BL21 (DE3) cells carrying pET22b-cnoX and pET22b-cnoX_C38A/C63A_ as follows. After an overnight preculture, cells were diluted in 500 mL of LB containing 100 mg/mL ampicillin and grown at 37°C. At the mid-log phase, we induced protein expression for 1 h with 1 mM IPTG. Cultures were cooled on ice for 10 min, and iodoacetamide (25 mM) was added. After 30 min, the cells were harvested, resuspended in 20 mL of buffer E, and sonicated. After centrifugation at 12,000*g* at 4°C, we added 150 µL of undiluted rabbit anti-GroEL antibody and 1 mL of protein A/G magnetic beads (Pierce) to the supernatant. After 30 min on a wheel at room temperature, magnetic beads were collected and resuspended with 1 mL of buffer E. After three washes with 1 mL of buffer E, GroEL was eluted with 100 µL of 0.1 M glycine pH 2.5.

**Atomic force microscopy**

Gold-coated glass coverslips and cantilevers (OMCL-TR4, Olympus Ltd.; nominal spring constant ∼0.02 N m^−1^) were incubated with 0.1 mg ml^−1^ of CnoX_N-link/C38A/C63A_ or GroEL_D490C_ solutions for 1 h, rinsed with buffer D, and then immediately used without dewetting.

For single-molecule force spectroscopy experiments, we prepared CnoX_N-link/C38A/C63A_ or GroEL_D490C_-functionalized surfaces and functionalized cantilevers as described above. We performed measurements at room temperature in 50 mM Tris pH 8, 150 mM NaCl, 1 mM EDTA with a Force Robot 300 AFM (JPK Instruments). Multiple (32 × 32) force–distance curves were recorded on areas of 500x500 nm^2^ with an applied force of 250 pN, a constant approach, and a retraction speed of 1,000 nm s^−1^. The histogram was generated by considering the force of the last rupture event for each curve. The spring constants of the cantilevers were measured by the thermal noise method. We analyzed these data with data-processing software from JPK Instruments.

**Native PAGE**

We performed Blue native electrophoresis analysis of the concentrated complex on a 3%–12% Bis-Tris gel (Life Technologies) following the manufacturer’s instructions. We identified the protein complex bands separated in the native electrophoresis via SDS-PAGE. Briefly, bands of interest were excised, boiled in SDS-PAGE sample buffer, and applied to the top of a polyacrylamide gel.

**Immunoblotting**

Samples were boiled before being loaded onto a precast NuPAGE Bis-Tris 12% gel (Life Technologies). We performed immunoblotting according to standard procedures using 1:5000 a-CnoX (rabbit serum, CER Groupe, Belgium) followed by a horseradish peroxidase-conjugated a-rabbit antibody (Sigma). We conducted chemiluminescence imaging (ECL Prime Western Blotting Detection Reagents; GE Healthcare) with an ImageQuant LAS 500 Camera (GE Healthcare Life Sciences).

**Finding mammalian homologs of CnoX**

The CnoX structure (<http://pfam.xfam.org/protein/P77395>) is organized in three domains, corresponding to PFAM entries *Thioredoxin* (<http://pfam.xfam.org/family/Thioredoxin>), *TPR_19* (<http://pfam.xfam.org/family/TPR_19>) and *TPR_20* (<http://pfam.xfam.org/family/TPR_20>). As of March 9^th^, 2022, these PFAMs contain 40,361 (*Thioredoxin*), 10,056 (*TPR_19*) and 44 (*TPR_20*) eukaryotic sequences, respectively. All 44 eukaryotic sequences were downloaded from UniProt (Consortium, 2021) and the corresponding AlphaFold three-dimensional models were built using ColabFold (Abadi et al., 2016). Most of these models contained all three domains *Thioredoxin*, *TPR_19* and *TPR_20*. Three representative structures are shown in Figure S6.

### Protein identification by MS

After in-gel digestion with trypsin, peptides were dissolved in solvent A (0.1% trifluoroacetic acid in 2% acetonitrile [ACN]), directly loaded onto a reversed-phase precolumn (Acclaim PepMap 100, Thermo Fisher Scientific), and eluted in backflush mode. We performed peptide separation using a reversed-phase analytical column (Acclaim PepMap RSLC, 0.075 x 250 mm, Thermo Fisher Scientific) with a linear gradient of 4%–27.5% solvent B (0.1% formic acid in 98% ACN) for 40 min, 27.5%–50% solvent B for 20 min, 50%–95% solvent B for 10 min, and holding at 95% for the last 10 min at a constant flow rate of 300 nL/min on an Ultimate 3000 RSLC system. We analyzed the peptides by an Orbitrap Fusion Lumos tribrid MS (Thermo Fisher Scientific). The peptides were subjected to a nanospray ionization source followed by MS/MS in a Fusion Lumos coupled online to a nano-LC. Intact peptides were detected in the Orbitrap at a resolution of 120,000. We selected peptides for MS/MS using a higher-energy collision dissociation setting of 30 and detected ion fragments in the Orbitrap at a resolution of 30,000. A data-dependent procedure that alternated between one MS scan and one MS/MS scan was applied for 3 s for ions above a threshold ion count of 2.0E4 in the MS survey scan with 40.0-s dynamic exclusion. An electrospray voltage of 2.1 kV was applied. We obtained MS1 spectra with an automatic gain control (AGC) target of 4E5 ions and a maximum injection time of 50 ms and MS2 spectra with an AGC target of 5E4 ions and a maximum injection set to dynamic. For MS scans, the m/z scan range was 375–1800. We processed the resulting MS/MS data using the Sequest HT search engine within Proteome Discoverer 2.4 SP1 against an *E. coli K12* protein database obtained from Uniprot (4,349 entries) Trypsin was specified as a cleavage enzyme allowing up to two missed cleavages, four modifications per peptide, and up to five charges. The mass error was set to 10 ppm for precursor ions and 0.1 Da for fragment ions. Oxidation on Met (+15.995 Da) and conversion of Gln (-17.027 Da) or Glu (-18.011 Da) to pyro-Glu at the peptide N-term were considered as variable modifications. We assessed the false discovery rate using Percolator, with the thresholds for proteins, peptides, and modification sites specified at 1%. For abundance comparison, we calculated abundance ratios by label-free quantification of the precursor intensities within Proteome Discoverer 2.4 SP1.

**Single-particle cryoEM imaging and data processing**

Prior to vitrification, we assessed the sample quality using negative-stain EM. For this step, 3 μL GroEL-CnoX at 0.4 mg/mL was applied to a glow-discharged formvar Cu400 grid (Electron Microscopy Sciences) and screened using in-house 120-kV JEOL JEM 1400 and 1400+ microscopes equipped with an LaB_6_ filament at the VIB-VUB facility for Bio Electron Cryogenic Microscopy (BECM). We vitrified the GroEL-CnoX sample in liquid ethane using a CP3 Cryoplunge (Gatan) plunger set to 100% humidity and room temperature. R2/1 (Quantifoil) grids were coated with graphene oxide (Sigma), and a 3-μL sample at 0.4 mg/mL was applied and manually back blotted for 4 s before plunging.

We collected data on an in-house (BECM) 300-kV JEOL CRYOARM 300 equipped with a K3 direct electron detector (Gatan). We collected 3,015 movies in counting mode, with a dose of 68.27 e/Å^2^ spread over 61 frames at a nominal magnification of 60.000x, corresponding to a pixel size of 0.784 Å. Images were motion-corrected via MotionCor2.1 (Zheng et al., 2017) using 5x5 patches with dose-weighting, and the contrast transfer function was estimated using ctffind4.1 (Rohou and Grigorieff, 2015). We picked particles using crYOLO 1.5 (Wagner et al., 2019) with a dataset-specific trained crYOLO network on six micrographs (709 picked particles). The picking quality and CnoX occupancy were assessed using ISAC2 (Yang et al., 2012) as implemented in the SPHIRE package (Moriya et al., 2017).

We performed all subsequent processing using RELION3.0. A total of 670,080 picked particles were extracted with a box size of 400 and were binned twice for processing. A c1 initial 3D model was generated *ab initio*. In all subsequent steps, we applied c7 symmetry throughout. After extensive rounds of 3D classification to optimize the density for CnoX on GroEL, we utilized 170,458 particles for the final reconstruction at 3.4 Å.

Final maps were density-modified using phenix.resolve (Terwilliger et al., 2020), and the model was fitted and symmetry-expanded in UCSF Chimera (Pettersen et al., 2004) using the second TPR domain of CnoX (helices 2_A_-2_A_’-2_B-_2_B_’-2_C_ [197-284] from PDB: 3QOU) and GroEL (pdb:1XCK) as starting coordinates. We conducted further docking and refinement using phenix.real.space (Afonine et al., 2018) with manual curation in COOT (Emsley and Cowtan, 2004). After the first refinement, we generated a second local resolution-sharpened map using LocScale (Jakobi et al., 2017), as implemented in the CCPEM project package (Burnley et al., 2017). We used this map together with the density-modified map to aid further map building and refinement in Phenix (Liebschner et al., 2019). The reported FSC-model curve, resolution, and local-resolution maps were calculated using the Phenix validation output and the Phenix implementation of local-resolution assessment.

**Table S1: CryoEM statistics table.**

|  | GroEL:CnoX  (EMDB-14352)  (PDB 7YWY) |
| --- | --- |
| **Data collection and processing** | JEOL CryoARM300, BECM, Brussels |
| Magnification | 60.000 |
| Voltage (kV) | 300 |
| Electron exposure (e–/Å^2^) | 68.3 |
| Defocus range (μm) | -0.5 to -3 |
| Pixel size (Å) | 0.784 |
| Symmetry imposed | C7 |
| Initial particle images (no.) | 670080 |
| Final particle images (no.) | 170458 |
| Map resolution (Å)  FSC threshold | 3.4  0.143 |
| Map resolution range (Å) | 3.1-7.6 |
| **Refinement** |  |
| Initial model used (PDB code) | 1XCK (GroEL) 3QOU (CnoX) |
| Model resolution (Å)  FSC threshold | 3.5 0.143 |
| Model resolution range (Å) |  |
| Map sharpening *B* factor (Å^2^) | Local Sharpening- Density Modification |
| Model composition  Non-hydrogen atoms  Protein residues  Ligands | 63994 8568 0 |
| *B* factors (Å^2^)  Protein  Ligand | 120.42 0 |
| R.m.s. deviations  Bond lengths (Å)  Bond angles (°) | 0.010 1.249 |
| Validation  MolProbity score  Clashscore  Poor rotamers (%) | 2.04  7.91 2.10 |
| Ramachandran plot  Favored (%)  Allowed (%)  Disallowed (%) | 94.75 4.76 0.49 |

**Table S2: Proteins involved in the mixed-disulfides with CnoX.**

Proteins that were pulled-down with the GroEL-CnoX complexes using a-GroEL antibodies and identified using LC-MS/MS. They were not detected when the experiments were repeated in cells expressing CnoX_no_cys_.

| **Accession**  **Number** | **Protein** | **Gene** | **GroEL**  **Substrates**  **(Kerner et al., 2005)** | **obligate substrates**  **(Fujiwara et al., 2010)** |
| --- | --- | --- | --- | --- |
| [P77357](https://www.uniprot.org/uniprot/P77357) | p-aminobenzoyl-glutamate hydrolase subunit A | *abgA* |  |  |
| P0AFG8 | Pyruvate dehydrogenase E1 component, PDH E1 component | *aceE* |  |  |
| P0A6A3 | Acetate kinase | *ackA* | X |  |
| P36683 | Aconitate hydratase B | *acnB* |  |  |
| P0A959 | Glutamate-pyruvate aminotransferase AlaA | *alaA* |  |  |
| P0A6C5 | Amino-acid acetyltransferase | *argA* |  |  |
| P23908 | Acetylornithine deacetylase | *argE* | X | X |
| P11875 | Arginine--tRNA ligase | *argS* |  |  |
| P77398 | Bifunctional polymyxin resistance protein ArnA | *arnA* |  |  |
| P0A8M0 | Asparagine--tRNA ligase | *asnS* |  |  |
| P0A6H1 | ATP-dependent Clp protease ATP-binding subunit ClpX | *clpX* |  |  |
| P00936 | Adenylate cyclase | *cyaA* |  |  |
| P23886 | ATP-binding/permease protein CydC | *cydC* |  |  |
| [P0ABI8](https://www.uniprot.org/uniprot/P0ABI8) | Cytochrome bo(3) ubiquinol oxidase subunit 1 | *cyoB* |  |  |
| P17846 | Sulfite reductase [NADPH] hemoprotein beta-component | *cysI* |  |  |
| P23845 | Sulfate adenylyltransferase subunit 1 | *cysN* |  |  |
| P0A6L2 | 4-hydroxy-tetrahydrodipicolinate synthase | *dapA* | X | X |
| P0A6P5 | GTPase Der der | *der* |  |  |
| P0ABQ0 | Coenzyme A biosynthesis bifunctional protein CoaBC | *dfp* |  |  |
| [P03004](https://www.uniprot.org/uniprot/P03004) | Chromosomal replication initiator protein DnaA | *dnaA* |  |  |
| P06710 | DNA polymerase III subunit tau | *dnaX* |  |  |
| P0A6P9 | Enolase | *eno* | X |  |
| P30845 | Phosphoethanolamine transferase EptA | *eptA* |  |  |
| P0CB39 | Phosphoethanolamine transferase EptC | *eptC* |  |  |
| P0AAI5 | 3-oxoacyl-[acyl-carrier-protein] synthase 2 | *fabF* | X | X |
| P32176 | Formate dehydrogenase-O major subunit | *fdoG* |  |  |
| P13036 | Fe(3+) dicitrate transport protein FecA | *fecA* |  |  |
| P46889 | DNA translocase FtsK | *ftsK* |  |  |
| P10121 | Signal recognition particle receptor FtsY | *ftsY* |  |  |
| P0A6M8 | Elongation factor G | *fusA* |  |  |
| P00370 | NADP-specific glutamate dehydrogenase | *gdhA* | X | X |
| P31120 | Phosphoglucosamine mutase | *glmM* |  |  |
| P0AEW6 | Guanosine-inosine kinase | *gsk* |  |  |
| P0ADG7 | Inosine-5'-monophosphate dehydrogenase | *guaB* |  |  |
| P0AES6 | DNA gyrase subunit B | *gyrB* |  |  |
| P0ACB2 | Delta-aminolevulinic acid dehydratase | *hemB* | X | X |
| P09127 | Protein HemX | *hemX* |  |  |
| [P25519](https://www.uniprot.org/uniprot/P25519) | GTPase HflX | *hflX* |  |  |
| P76658 | Bifunctional protein HldE | *hldE* |  |  |
| P08957 | Type I restriction enzyme EcoKI M protein | *hsdM* |  |  |
| P69741 | Hydrogenase-2 small chain | *hybO* |  |  |
| P08200 | Isocitrate dehydrogenase [NADP] | *icd* |  |  |
| [P04968](https://www.uniprot.org/uniprot/P04968) | L-threonine dehydratase biosynthetic IlvA | *ilvA* |  |  |
| P0A705 | Translation initiation factor IF-2 | *infB* |  |  |
| P0A6B7 | Cysteine desulfurase IscS | *iscS* | X |  |
| P00803 | Signal peptidase I, SPase I | *lepB* |  |  |
| [P32099](https://www.uniprot.org/uniprot/P32099) | Lipoate-protein ligase A | *lplA* |  |  |
| P45464 | Penicillin-binding protein activator LpoA | *lpoA* |  |  |
| P08660 | Lysine-sensitive aspartokinase 3 | *lysC* |  |  |
| P15977 | 4-alpha-glucanotransferase | *malQ* |  |  |
| P21517 | Maltodextrin glucosidase | *malZ* |  |  |
| P07623 | Homoserine O-succinyltransferase | *metA* |  |  |
| P25665 | 5-methyltetrahydropteroyltriglutamate--homocysteine methyltransferase | *metE* |  |  |
| P13009 | Methionine synthase | *metH* |  |  |
| P0A817 | S-adenosylmethionine synthase | *metK* | X | X |
| P30750 | Methionine import ATP-binding protein MetN | *metN* |  |  |
| [P30958](https://www.uniprot.org/uniprot/P30958) | Transcription-repair-coupling factor | *mfd* |  |  |
| P0ABB8 | Magnesium-transporting ATPase | *mgtA* |  |  |
| P63386 | Intermembrane phospholipid transport system ATP-binding protein MlaF | *mlaF* |  |  |
| P25522 | tRNA modification GTPase MnmE | *mnmE* |  |  |
| P0AD65 | Peptidoglycan D,D-transpeptidase MrdA | *mrdA* |  |  |
| P60752 | ATP-dependent lipid A-core flippase | *msbA* |  |  |
| P60293 | Chromosome partition protein MukF | *mukF* |  |  |
| P17952 | UDP-N-acetylmuramate--L-alanine ligase | *murC* |  |  |
| P22188 | UDP-N-acetylmuramoyl-L-alanyl-D-glutamate--2,6-diaminopimelate ligase | *murE* |  |  |
| P22634 | Glutamate racemase | *murI* |  |  |
| P00452 | Ribonucleoside-diphosphate reductase 1 subunit alpha | *nrdA* |  |  |
| P31979 | NADH-quinone oxidoreductase subunit F | *nuoF* | X |  |
| P0A910 | Outer membrane protein A | *ompA* | X |  |
| P02931 | Outer membrane porin F | *ompF* | X |  |
| P0ABF1 | Poly(A) polymerase I | *pcnB* | X |  |
| [P0AFL6](https://www.uniprot.org/uniprot/P0AFL6) | Exopolyphosphatase, ExopolyPase | *ppx* |  |  |
| P07012 | Peptide chain release factor RF2 | *prfB* |  |  |
| [P0A7B5](https://www.uniprot.org/uniprot/P0A7B5) | Glutamate 5-kinase | *proB* |  |  |
| P14175 | Glycine betaine/proline betaine transport system ATP-binding protein ProV | *proV* |  |  |
| P0A7D4 | Adenylosuccinate synthetase | *purA* | X |  |
| P0AB89 | Adenylosuccinate lyase | *purB* |  |  |
| P09546 | Bifunctional protein PutA | *putA* |  |  |
| P24554 | DNA repair protein RadA | *radA* |  |  |
| P60240 | RNA polymerase-associated protein RapA | *rapA* |  |  |
| P0AAZ4 | Replication-associated recombination protein A | *rarA* |  |  |
| P04993 | RecBCD enzyme subunit RecD | *recD* |  |  |
| [P24230](https://www.uniprot.org/uniprot/P24230) | ATP-dependent DNA helicase RecG | *recG* |  |  |
| [P27833](https://www.uniprot.org/uniprot/P27833) | dTDP-4-amino-4,6-dideoxygalactose transaminase | *rffA* |  |  |
| P25888 | ATP-dependent RNA helicase RhlE | *rhlE* | X |  |
| P0AG30 | Transcription termination factor Rho | *rho* | X |  |
| P36979 | Dual-specificity RNA methyltransferase RlmN | *rlmN* |  |  |
| P09155 | Ribonuclease D | *rnd* |  |  |
| [P21513](https://www.uniprot.org/uniprot/P21513) | Ribonuclease E | *rne* |  |  |
| P62399 | 50S ribosomal protein L5 | *rplE* |  |  |
| P0A8T7 | DNA-directed RNA polymerase subunit beta | *rpoC* |  |  |
| P0AG67 | 30S ribosomal protein S1 | *rpsA* |  |  |
| P0A7V0 | 30S ribosomal protein S2 | *rpsB* |  |  |
| P0A7V8 | 30S ribosomal protein S4 | *rpsD* |  |  |
| P06992 | Ribosomal RNA small subunit methyltransferase A | *rsmA* |  |  |
| [P77611](https://www.uniprot.org/uniprot/P77611) | Ion-translocating oxidoreductase complex subunit C | *rsxC* |  |  |
| P16095 | L-serine dehydratase 1 | *sdaA* |  |  |
| P30744 | L-serine dehydratase 2 | *sdaB* |  |  |
| P0AC41 | Succinate dehydrogenase flavoprotein subunit | *sdhA* | X | X |
| P0AG90 | Protein translocase subunit SecD | *secD* |  |  |
| P0AGA2 | Protein translocase subunit SecY | *secY* |  |  |
| P23721 | Phosphoserine aminotransferase | *serC* | X | X |
| P00934 | Threonine synthase | *thrC* |  |  |
| P0AGG8 | Metalloprotease TldD | *tldD* | X | X |
| P0A855 | Tol-Pal system protein TolB | *tolB* |  |  |
| P14294 | DNA topoisomerase 3 | *topB* |  |  |
| P0AGI8 | Trk system potassium uptake protein TrkA | *trkA* |  |  |
| P23003 | tRNA/tmRNA (uracil-C(5))-methyltransferase | *trmA* | X |  |
| P0CE48 | Elongation factor Tu 2 | *tufB* |  |  |
| P0DTT0 | 50S ribosomal subunit assembly factor BipA | *typA* | X |  |
| P07604 | Transcriptional regulatory protein TyrR | *tyrR* |  |  |
| P0AGJ9 | Tyrosine--tRNA ligase | *tyrS* |  |  |
| P76373 | UDP-glucose 6-dehydrogenase | *ugd* |  |  |
| P0A698 | UvrABC system protein A | *uvrA* |  |  |
| P75829 | Uncharacterized protein YbjX | *ybjX* |  |  |
| P75863 | Uncharacterized protein YcbX | *ycbX* |  |  |
| P75867 | Putative Lon protease homolog | *ycbZ* |  |  |
| P24188 | tRNA uridine(34) hydroxylase | *yceA* |  |  |
| P77748 | Uncharacterized protein YdiJ | *ydiJ* |  |  |
| P76272 | Intermembrane transport protein YebT | *yebT* |  |  |
| P36928 | Uncharacterized chaperone protein YegD | *yegD* | X |  |
| P76403 | tRNA hydroxylation protein P | *yegQ* |  |  |
| P0ADR8 | Pyrimidine/purine nucleotide 5'-monophosphate nucleosidase | *ygdH* |  |  |
| Q46861 | UPF0313 protein YgiQ | *ygiQ* |  |  |
| P64612 | Cell division protein ZapE | *yhcM* |  |  |
| P46837 | Protein YhgF | *yhgF* |  |  |
| P77173 | Cell division protein ZipA | *zipA* |  |  |

**Table S3: Bacterial strains used in this study.**

| **Strain** | **Genotype** | **Plasmid** | **Source and notes** |
| --- | --- | --- | --- |
| JF179 | MG1655 *wt* |  | From Jim Bardwell (U of Michigan) |
| ED002 | MG1655 *ΔcnoX* |  | Goemans et al., 2018 |
| ED004 | MG1655 *ΔcnoX* | pET22b-cnoX | Goemans et al., 2018 |
| ED006 | MG1655 *ΔcnoX* | pET22b-cnoX_C-His_ | This study |
| ED005 | MG1655 *ΔcnoX* | pET22b-cnoX_ΔCter_ | This study |
| CG233 | BL21 (DE3) | pET22b-groL | Goemans et al., 2018 |
| ED035 | BL21 (DE3) | pET22b-groL_D490C_ | This study |
| ED126 | BL21 (DE3) | pET22b-groL_R452A/E461A/S463A/V464A_ | This study; Weissman et al., 1995, 1996 |
| ED187 | BL21 (DE3) | pET22b-groL_G298A/T299L/V300K/E304L/I305K/M307K/R345L_ | This study |
| CG183 | BL21 (DE3) | pET22b-groS | This study |
| CG232 | BL21 (DE3) | pET22b-cnoX | Goemans et al., 2018 |
| CG229 | BL21 (DE3) | pET22b-cnoX_ΔCter_ | This study |
| CG191 | BL21 (DE3) | pET22b-cnoX_C-His_ | This study |
| CG260 | BL21 (DE3) | pET22b-cnoX_C38A/C63A_ | Goemans et al., 2018 |
| ED042 | BL21 (DE3) | pET22b-cnoX_N-link/C38A/C63A_ | This study |
| ED145 | BL21 (DE3) | pET22b-mhsp60 | This study |
| ED049 | BL21 (DE3) | pACYCDuet-1-cnoX_N-Strep_ | This study |

**Table S4: Plasmids used in this study.**

| **Plasmid** | **Vector** | **Protein encoded** | | **Source and notes** | |
| --- | --- | --- | --- | --- | --- |
| pET22b(+) | IPTG-inducible P_lac_, ampicillin |  | | Novagen | |
| pET22b-cno*X* | pET22b(+) | CnoX | | Goemans et al., 2018 | |
| pET22b-groL | pET22b(+) | GroEL | | Goemans et al., 2018 | |
| pET22b-groL_D490C_ | pET22b(+) | GroEL_D490C_ | | This study | |
| pET22b-groL_R452A/E461A/S463A/V464A_ | pET22b(+) | GroEL_R452A/E461A/S463A/V464A_ | | This study; Weissman et al., 1995, 1996 | |
| pET22b-groL_G298A/T299L/V300K/E304L_ | pET22b(+) | GroEL_G298A/T299L/V300K/E304L/_ | | This study | |
| _/I305K/M307K/R345L_ |  | _I305K/M307K/R345L_ | |  | |
| pET22b-groS | pET22b(+) | GroES | | This study | |
| pET22b-cnoX_ΔCter_ | pET22b(+) | CnoX without the last 10 C-terminal amino acids | | This study | |
| pET22b-cnoX_C-His_ | pET22b(+) | CnoX fused with a C-terminal 6xHis tag | | This study | |
| pET22b-cnoX_C38A/C63A_ | pET22b(+) | CnoX_C38A/C63A_ | | Goemans et al., 2018 | |
| pET22b-cnoX_N-link/C38A/C63A_ | pET22b(+) | CnoX_C38A/C63A_ with an N-terminal cysteine linker (M/A/C/A/G residues) | | This study | |
| pACYCDuet-1 | IPTG-inducible P_lac_, chloramphenicol |  | | Novagen | |
| pACYCDuet-1-cnoX_N-Strep_ | pACYCDuet-1 | CnoX fused with an N-terminal  Strep-tag | | This study | |
| MAC-C-CH60 | MAC-tag-C | CH60 from *H. sapiens* (HSPD1) | | Addgene | |
| pET22b-mhsp60 | pET22b(+) | CH60 with the MTS substituted by M/G/S residues | | This study | |
| pET15b-cnoX | pET15b | | CnoX with thrombin-cleavable N-terminal 6xHis tag | | Lin & Wilson, 2011 |
| pET15b-groS | pET15b | | GroES with thrombin-cleavable N-terminal 6xHis tag | | Lin & Wilson, 2011 |
| pET15b-groL | pET15b | | Native GroEL with no tag | | This study |

**Table S5: Primers used in this study.**

| **Primer** | **Sequence (5' -> 3')** | **Comments** |
| --- | --- | --- |
| AGo230_cnoX_Strep_NcoI_F | AAAACCATGGCAAGCTGGAGCCACCCGCAGTTCGAAAAGTCCGTAGAAAATATTGTCAAC | cloning (pACYCDuet-1-cnoX_N-Strep_) |
| AGo231_cnoX_Strep_BamHI_R | AAAAGGATCCTCAATACAACAATGCATACAGC | cloning (pACYCDuet-1-cnoX_NterStrep_) |
| CGo174_GroES_NdeI_F | GAGCATATGAAGTTTCGTCCC | cloning (pET22b-groS) |
| CGo185_GroES_stop_XhoI_R | GAGCTCGAGTTAGGCTTCGACCACGCC | cloning (pET22b-groS) |
| CGo178_cnoX_H6_NdeI_F | GAGCATATG*CACCACCACCACCACCAC*TCCCTGATCGGC | cloning (pET22b-cnoX_C-His_) |
| CGo71_cnoX_H6-stop_XhoI_R | CTCCTCGAGTCAGTGGTGGTGGTGGTGGTGGCTGAAGAGGATCGAGG | cloning (pET22b-cnoX_C-His_) |
| CGo57_cnoX_NdeI_F | GAGCATATGATGTCCCTGATCGGC | cloning (pET22b-cnoX_ΔCter_) |
| CGo234_cnoXdel10_HindIII_stop_R | CTCAAGCTTTCACTTCGACGCCAGTGCATCACC | cloning (pET22b-cnoX_ΔCter_) |
| EDo227_22b(mHSP60)_F | TGAGATCCGGCTGCTAAC | cloning (pET22b-mhsp60) |
| EDo228_22b(mHSP60)_R | ATGTATATCTCCTTCTTAAAGTTAAACAAAATTATTTC | cloning (pET22b-mhsp60) |
| EDo229_(22b)mHSP60_F | TTTAAGAAGGAGATATACATATGGGAAGTGCCAAAGATGTAAAATTTGGTGC | cloning (pET22b-mhsp60) |
| EDo230_(22b)mHSP60_R | TTGTTAGCAGCCGGATCTCAGAACATGCCACCTCCCATAC | cloning (pET22b-mhsp60) |
| EDo31_GroEL-D490C_F | TGGATCCAGGATACCCATGCAGATCATGTTGCCGTATTCT | cloning (pET22b-groEL_D490C_) |
| EDo32_GroEL-D490C_R | AGAATACGGCAACATGATCTGCATGGGTATCCTGGATCCA | cloning (pET22b-groEL_D490C_) |
| EDo321_AD_groEL_G298A-T299L-E304L-R345L_Gibson_frag1_F | GACTGGCGCGCTTGTGATCTCTGAACTTATCG | cloning (pET22b-groL_G298A/T299L/V300K/E304L/I305K/M307K/R345L_) |
| EDo322_AD_groEL_G298A-T299L-E304L-R345L_Gibson_frag1_R | GTCACGCTCGTCGTTTGGTATGGCTTCATTC | cloning (pET22b-groL_G298A/T299L/V300K/E304L/I305K/M307K/R345L_) |
| EDo323_AD_groEL_G298A-T299L-E304L-R345L_Gibson_frag2_F | TACCAAACGACGAGCGTGACACCACGATG | cloning (pET22b-groL_G298A/T299L/V300K/E304L/I305K/M307K/R345L_) |
| EDo324_AD_groEL_G298A-T299L-E304L-R345L_Gibson_frag2_R | AGATCACAAGCGCGCCAGTCAGGGTTGCGATATC | cloning (pET22b-groL_G298A/T299L/V300K/E304L/I305K/M307K/R345L_) |
| EDo325_AD_groEL_G298A-T299L-V300K-E304L-I305K-M307K-R345L_Gibson_frag1_F | AATCCAGGGCCTTGTTGCTCAGATCCGTCAGCAG | cloning (pET22b-groL_G298A/T299L/V300K/E304L/I305K/M307K/R345L_) |
| EDo326_AD_groEL_G298A-T299L-V300K-E304L-I305K-M307K-R345L_Gibson_frag1_R | CTTTACCTTTAAGTTCAGAGATTTTAAGCGCGCCAGTCAGGGT | cloning (pET22b-groL_G298A/T299L/V300K/E304L/I305K/M307K/R345L_) |
| EDo327_AD_groEL_G298A-T299L-V300K-E304L-I305K-M307K-R345L_Gibson_frag2_F | AAAATCTCTGAACTTAAAGGTAAAGAGCTGGAAAAAGC | cloning (pET22b-groL_G298A/T299L/V300K/E304L/I305K/M307K/R345L_) |
| EDo328_AD_groEL_G298A-T299L-V300K-E304L-I305K-M307K-R345L_Gibson_frag2_R | GAGCAACAAGGCCCTGGATTGCAGCTTC | cloning (pET22b-groL_G298A/T299L/V300K/E304L/I305K/M307K/R345L_) |
| EDo34_22b_MACAG_cnoX_F | ATGGCTTGTGCTGGTATGTCCGTAGAA | cloning (pET22b- cnoX_N-link/C38A/C63A_) |
| EDo35_22b_MACAG_cnoX_R | ACCAGCACAAGCCATCATATGTATAATC | cloning (pET22b-cnoX_N-link/C38A/C63A_) |
| EDo92_22b_groEL_R452A_F | AGCTCCGCTGGCTCAGATCGTATTGAACTGC | cloning (pET22b-groL_R452A/E461A/S463A/V464A_) |
| EDo93_22b_groEL_R452A_R | TCCATTGCACGCAGTGCA | cloning (pET22b-groL_R452A/E461A/S463A/V464A_) |
| EDo94_22b_groEl_E461A/S463A/V464A_F | GCTGCTGTTGCTAACACCGTTAAAG | cloning (pET22b-groL_R452A/E461A/S463A/V464A_) |
| EDo95_22b_groEl_E461A/S463A/V464A_R | CGGAGCTTCGCCGCAGTTCAATAC | cloning (pET22b_groL_R452A/E461A/S463A/V464A_) |
| CnoX_NdeI_for | GCCGACGCCCCTTGCATATGTCCGTAGAAAATATTGTC | cloning (pET15b-cnoX) |
| CnoX_XhoI_rev | CCGTGCTTTCTTGCTCGAGTCAATACAACAATGCATACAGC | cloning (pET15b-cnoX) |
| GroES_NdeI_for | CCTGAGAAGCGTTCCATATGAATATTCGTCCATTGCATGATCG | cloning (pET15b -groS) |
| GroES_XhoI_rev | CCGTGCTGCCTTGCTCGAGTTACGCTTCAACAATTGCCAGAATG | cloning (pET15b -groS) |
| GroEL_mHis_for | CTTTAAGAAGGAGATATACCCATATGGCAGCTAAAGACGTAAAATTC | cloning (pET15b-groL) |
| GroEL_mHis_rev | GAATTTTACGTCTTTAGCTGCCATATGGGTATATCTCCTTCTTAAAG | cloning (pET15b-groL) |

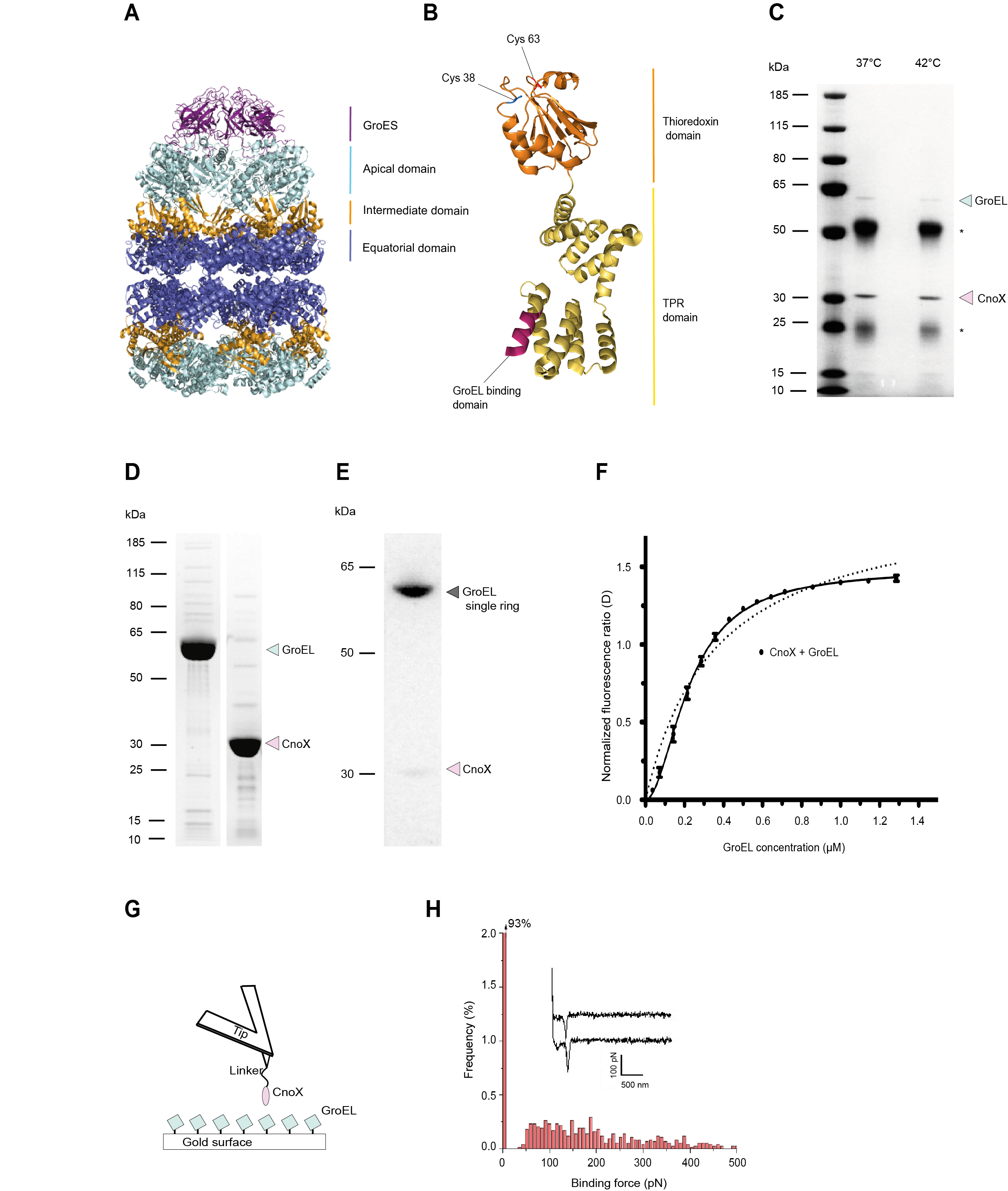

**Figure S1**

**(A)** Structure of the GroEL-GroES complex from *E. coli* (PDB: 1aon). The apical, hinge, and equatorial domains of GroEL are shown in light cyan, orange, and slate, respectively. GroES is shown in deep purple.

**(B)** Structure of *E. coli* CnoX (PDB: 3QOU) (Lin and Wilson, 2011). The N-terminal thioredoxin domain, with its two cysteine residues, is shown in orange. The C-terminal TPR domain is shown in gold. The last C-terminal residues are shown in purple.

**(C)** The amount of GroEL that co-elutes with CnoX does not increase when CnoX is pulled-down from extracts prepared from cells exposed to 42°C for 1 h instead of 37°C. We pulled-down CnoX using α-CnoX antibodies. The SDS-PAGE gel, stained with Coomassie blue, is representative of >3 replicates. * indicates the light and heavy chains of the antibodies.

**(D)** GroEL and CnoX were purified to near homogeneity.

**(E)** GroEL_R452A/E461A/S463A/V464A_, a GroEL variant that forms a single ring, co-elutes with CnoX from a size-exclusion chromatography column.

**(F)** Binding of FM-CnoX to GroEL was monitored via the fluorescence emission intensity of the fluorescein label on FM-CnoX. We performed measurements in triplicate, with the standard deviations indicated. The dotted line indicates the best fit for the noncooperative model, and the solid line presents a fitted curve for the positively cooperative model, with K_d_=227 nM and Hill coefficient (nH)=1.9.

**(G)** Schematic representation of the AFM setup for single-molecule force spectroscopy experiments: GroEL is attached to a gold-coated glass coverslip via a D490C substitution, and CnoX is attached to the cantilever via an N-terminal cysteine-containing linker.

**(H)** AFM measurement of the specific adhesion force. Two typical adhesion force curves are shown in the inset. We measured a specific binding force of 175±75 pN (mean and standard deviation out of a total of 5000 curves from three independent experiments). In these experiments, we used a GroEL mutant (GroEL_D490C_) with a cysteine residue exposed on its equatorial domain to allow binding to the gold surface and a CnoX variant (CnoX_N-link/C38A/C63A_) lacking the two cysteines of the thioredoxin domain­­ to prevent disulfide bond formation between CnoX and GroEL.

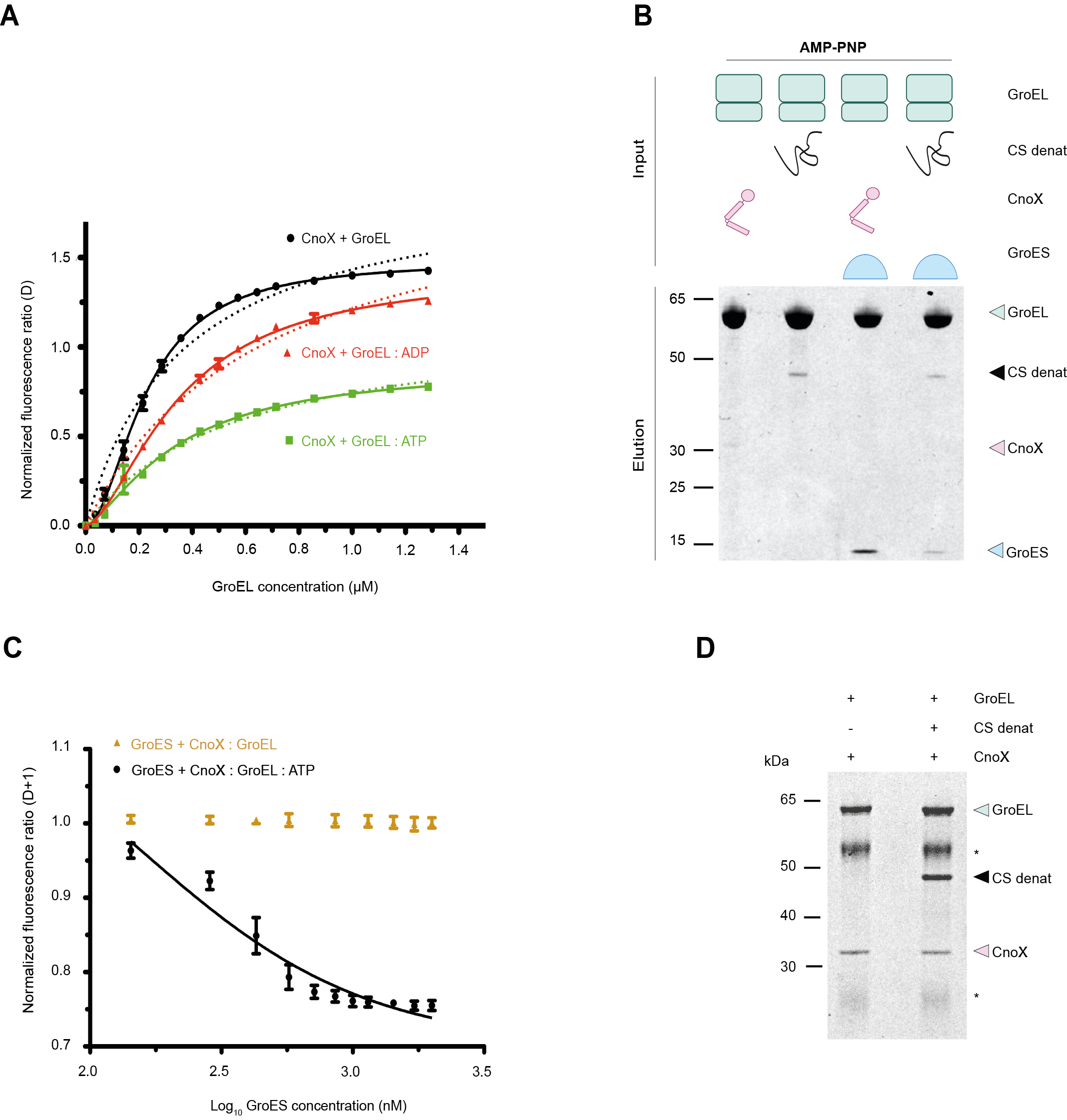

**Figure S2**

**(A)** Nucleotide binding to GroEL decreases the affinity for CnoX. We monitored the binding of FM-CnoX to GroEL in the presence of ADP using the fluorescence emission intensity of the fluorescein label on FM-CnoX. Dotted lines indicate best-fit curves for the noncooperative model, and solid lines present best-fit curves for the positively cooperative binding model. ADP occupancy on GroEL reduces the binding affinity of FM-CnoX to 350 nM. Error bars represent standard deviations from the results of three or more independent measurements.

**(B)** CnoX and unfolded CS co-elute with GroEL from a gel filtration column. Addition of GroES triggers the release of CnoX from GroEL, while CS remains bound to GroEL. We performed size-exclusion chromatography in the presence of 50 µM adenylyl-imidodiphosphate (AMP-PNP), a non-hydrolysable ATP analogue, and analyzed fractions by SDS-PAGE. The results are representative of >3 experiments.

**(C)** GroES displaces bound CnoX from GroEL. We added GroES to preformed FM-CnoX:GroEL in the absence (brown) and presence (black) of ATP and monitored the fluorescence intensity of FM-CnoX. The addition of GroES causes FM-CnoX:GroEL to dissociate with a K_i_ of 47 nM only when ATP is present. Error bars represent standard deviations from the results of three or more independent measurements.

**(D)** The GroEL-CnoX complex was incubated with or without unfolded CS. We pulled-down GroEL using specific α-GroEL antibodies. Both CnoX and unfolded CS coimmunoprecipitated with GroEL, indicating that CnoX binding does not prevent GroEL from recruiting unfolded CS. * indicates the light and heavy chains of the antibodies.
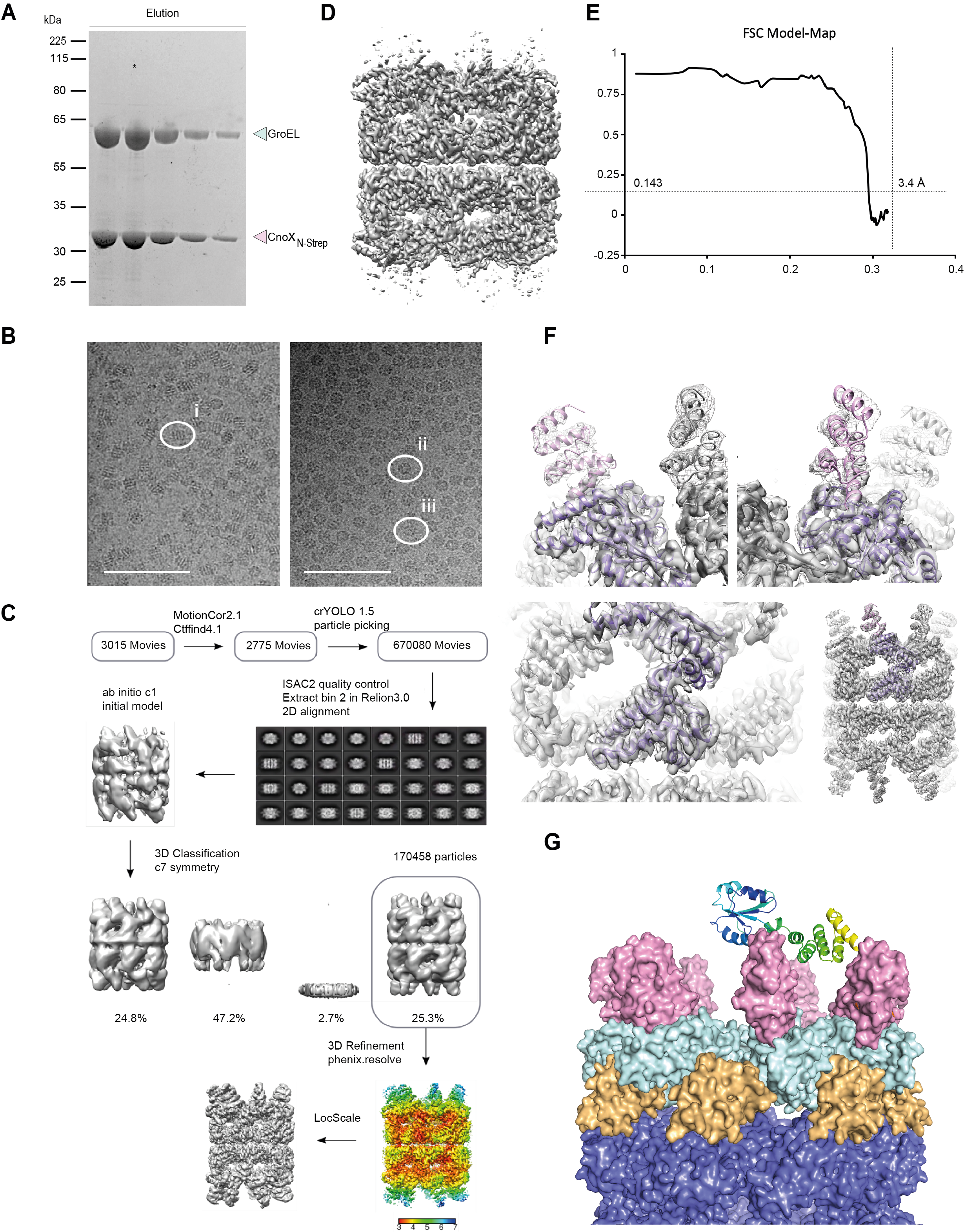

**Figure S3**

**(A)** Affinity purification of GroEL-CnoX (10:1 molar ratio). * indicates the fraction used for grid preparation.

**(B)** Representative cryoEM micrographs in thin (left) and thick (right) ice, displaying predominately side (i), top (ii), and tilted (iii) views (scale bar: 100 nm; i, ii, iii labeled as in Fig. 2B).

**(C)** Single-particle cryoEM processing pipeline. We optimized 3D map representations for CnoX density.

**(D)** Reconstructed 3D electron potential map showing good density for GroEL and sparse density for CnoX spokes.

**(E)** Fourier shell correlation (FSC) model map, as generated by Phenix.

**(F)** Density-fit illustrations of the GroEL:CnoX structure. The final 3D reconstruction is displayed as density-modified (surface) and locally sharpened (mesh) maps, which were best-suited for building GroEL and CnoX, respectively. The 3D reconstructions present a close-up view of the GroEL–CnoX contact region (top left and right, 90° rotated), a close-up view of the GroEL equatorial region (lower left), and a global particle view (lower right). Individual GroEL (purple) and CnoX (pink) subunits are highlighted for clarity.

**(G)** Side view of the GroEL-CnoX structure as resolved and built into the 3D cryoEM reconstruction (shown in surface representation) and the relative positioning of the CnoX thioredoxin domain (ribbon representation) based on a superimposition of the X-ray structure of *E. coli* CnoX (PDB: 3QOU) onto the CnoX TPR domain resolved in the 3D cryoEM reconstruction.

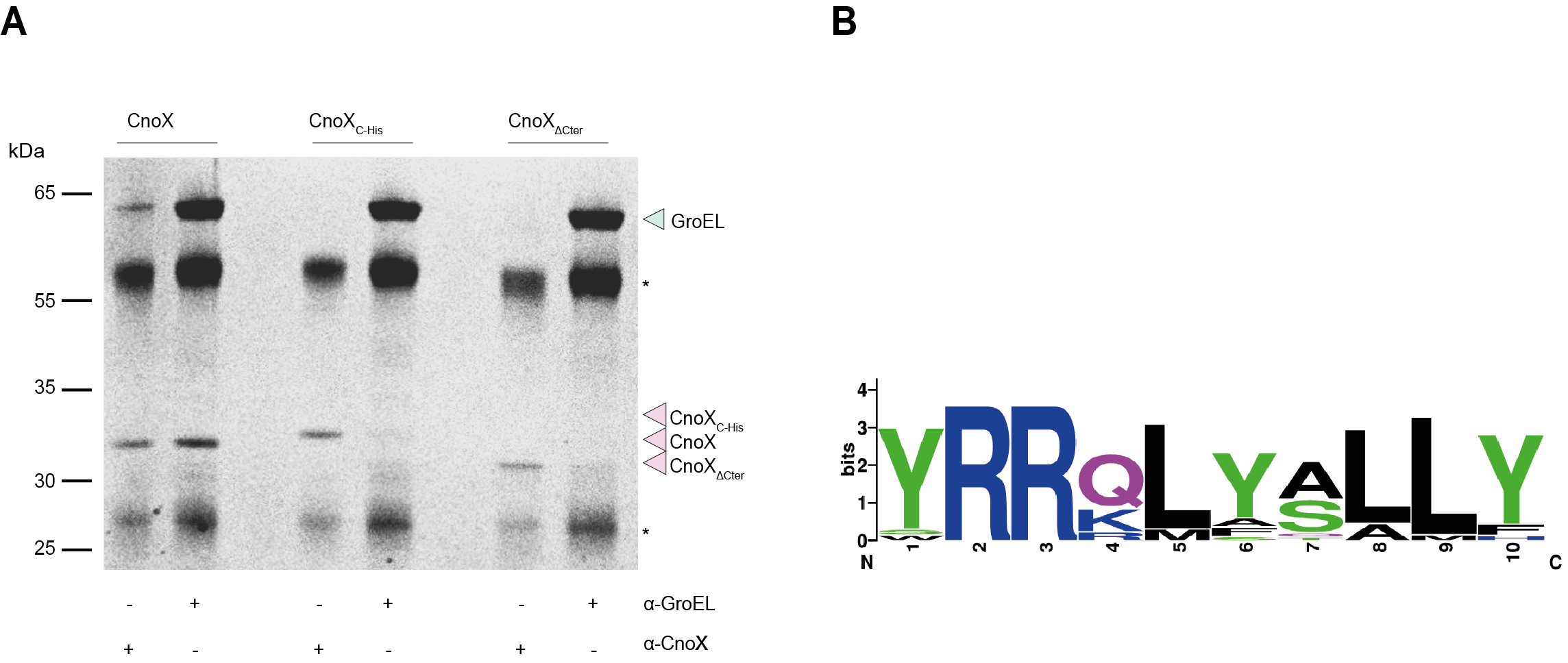

**Figure S4**

**(A)** GroEL was mixed with CnoX, CnoX_∆C-ter_, or CnoX_C-His_. We then used specific α-CnoX or α-GroEL antibodies to pull-down the corresponding protein. We observed an interaction between CnoX and GroEL only for wild-type CnoX, not for CnoX_∆C-ter_ or CnoX_C-His_, indicating that CnoX binds GroEL via its C-terminus. The SDS-PAGE gel, stained with Coomassie blue, is representative of >3 replicates. * indicates the light and heavy chains of the antibodies.

**(B)** Sequence logo showing the high conservation of the 10 C-terminal residues of CnoX among *E. coli, Shigella flexneri, Klebsiella pneumoniae, Enterobacter ludwigii, Serratia plymuthica, Pseudomonas fluorescens, Pantoea ananatis, Stenotrophomonas sp. DAIF1, Citrobacter freundii, Erwinia sp. Ejp617, Halomonas sp. HL-93, Cronobacter turicensis, Marinobacter sp. BSs20148, Salmonella enterica, Pseudomonas aeruginosa, Yersinia enterocolitica, Burkholderia gladioli,* and *Stèenotrophomonas maltophilia* (https://weblogo.berkeley.edu/).

**Figure S5**

Side-view surface representation of the cryoEM structure of GroEL–CnoX (left), side by side with structures of GroEL in different conformational states: the apo or T state (PDB: 1grl; (Braig *et al.*, 1994)), the Rs2 and Rd-open states (PDB: 4aar and 4ab3, respectively; (Clare *et al.*, 2012)), and the GroES-bound ES state (PDB: 1svt; (Chaudhry et al., 2004)). The binding paratope (shown in sea green) is accessible in the T, Rs, and Rs/Rd-open states, but becomes inaccessible in the GroES-bound ES state. In addition, sterically, the different conformational states of the GroEL apical domain would be compatible with CnoX binding, except for the ES state. CnoX is shown in pink. The GroEL equatorial, intermediate, and apical domains are shown in slate, orange, and light cyan. GroES is shown in fuchsia.

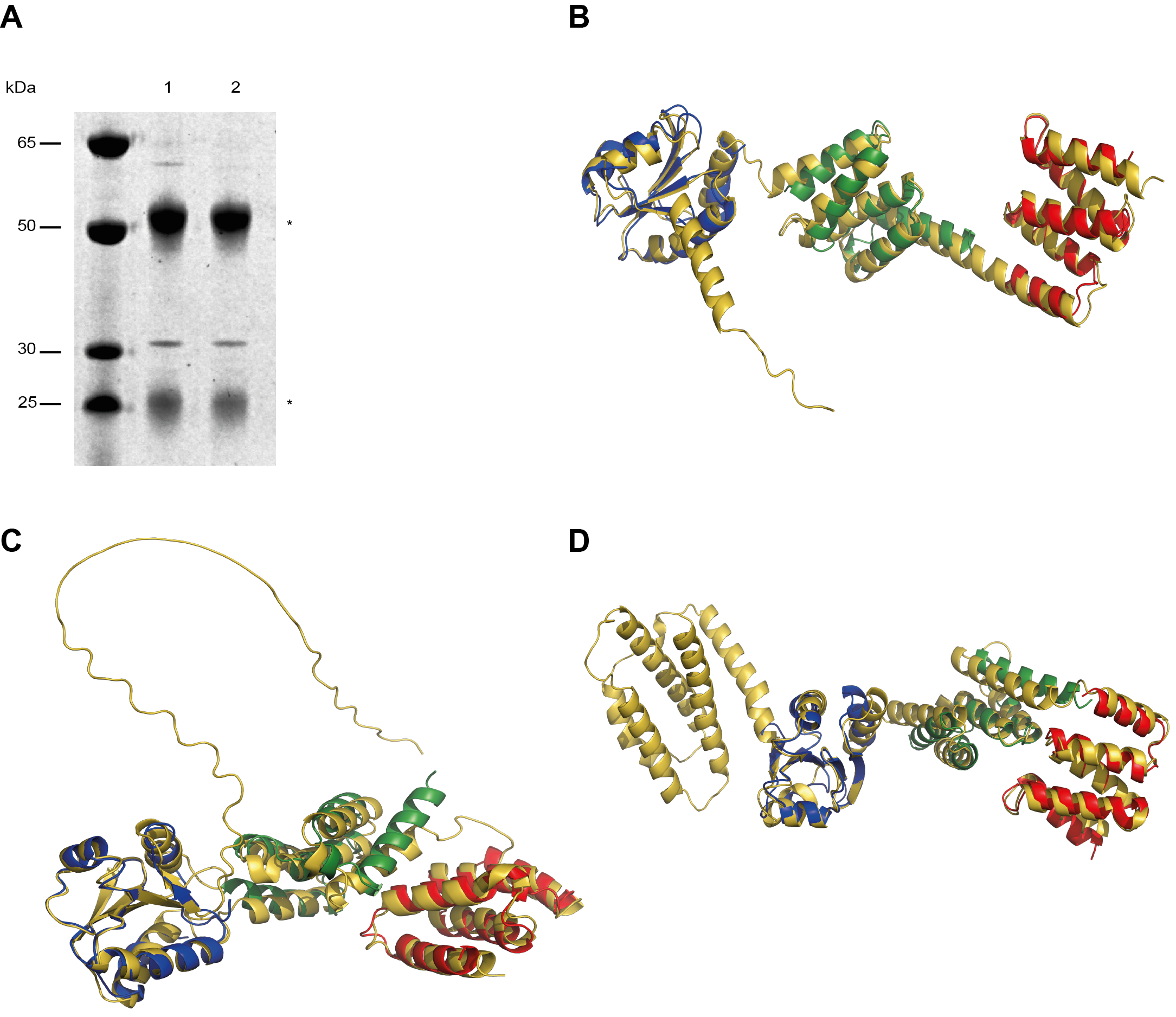

**Figure S6**

**(A)** GroEL (lane 1) or human mitochondrial Hsp60 (mHsp60, migrates slightly lower than GroEL; lane 2) co-elutes with CnoX when CnoX is pulled-down from cell extracts using α-CnoX antibodies. In these experiments, we expressed GroEL and mHsp60 in BL21 (DE3) cells. The SDS-PAGE gel, stained with Coomassie blue, is representative of >3 replicates. * indicates the light and heavy chains of the antibodies.

**(B,C,D)** Representative AlphaFold three-dimensional models of mammalian homologs of CnoX. b) *Stentor coeruleus* (UniProt A0A1R2CPV9), c) *Emiliania huxleyi* (UniProt R1DRS2) and d) *Pseudocohnilembus persalinus* (UniProt A0A0V0QL30). The domains *thioredoxin*, *TPR_19* and *TPR_20* (colored in blue, green and red, respectively) of CnoX (PDB code 3QOU) are superposed on these AlphaFold models (colored in gold).
